# Dendrite-wide voltage and Ca^2+^ imaging reveals structures of dendritic excitations during behavior

**DOI:** 10.64898/2026.01.18.700189

**Authors:** Xiang Wu, Byung Hun Lee, Pojeong Park, J. David Wong-Campos, Junjie Xu, Sarah E. Plutkis, Luke D. Lavis, Adam E. Cohen

**Affiliations:** Department of Chemistry and Chemical Biology, Harvard University, Cambridge, MA, USA; Department of Brain Sciences, DGIST, Daegu, Republic of Korea; Janelia Research Campus, Howard Hughes Medical Institute, Ashburn, Virginia, USA; Department of Physics, Harvard University, Cambridge, MA, USA

## Abstract

The dynamic interplay of dendritic voltage and Ca^2+^ shapes neuronal integration and synaptic plasticity, but the voltage-Ca^2+^ relationship during behavior remains poorly defined. Using dual- color Patterned Illumination Spinning disk COnfocal (PISCO) microscopy, we simultaneously mapped voltage and Ca^2+^ dynamics throughout the dendritic arbors of hippocampal CA2 pyramidal neurons during spatial navigation. Fast Na^+^ spikes usually initiated near the soma and back-propagated throughout the dendritic tree. Simple spikes primarily drove somatic and proximal Ca^2+^ increases, whereas complex spikes also engaged branch-selective apical Ca^2+^. In contrast, the voltage signals varied relatively smoothly across the dendritic tree. Dendrite-restricted excitations without concurrent somatic events were rare. We developed a biophysics-inspired model that accurately predicted local Ca^2+^ amplitude from local voltage waveforms across subcellular compartments and event types. Locomotion and spatial context shifted the relative prevalence of simple and complex spikes, but the same compact biophysical model captured the local voltage-Ca^2+^ relationships across behavioral states and contexts. We further found that local subthreshold depolarizations in apical dendrites preceding a plateau potential correlated with local Ca^2+^ amplitude during the plateau. Our results support a model in which conjunction of voltage back-propagation and branch-specific inputs together recruit regenerative dendritic electrogenesis and somatic bursting.

## Introduction

The dynamic interplay of membrane voltage and intracellular Ca^2+^ is the key electrical-chemical coupling step in the nervous system. The two signals are reciprocally coupled: membrane depolarization induces Ca^2+^ entry via voltage-gated Ca^2+^ channels (VGCCs) and N-methyl-D- aspartate (NMDA) receptors (with bound glutamate)^1,2^. This Ca^2+^ influx, in turn, contributes to dendritic electrogenesis. Ca^2+^-activated K⁺ channels^3,4^ and slower Ca^2+^-dependent signaling pathways^3^ also feed back onto voltage and excitability. Ca^2+^ signaling links dendritic activation to synaptic plasticity^5,6^, retrograde signaling^7^, and in some cases excitotoxicity^8^.

*In vivo*, dendritic Ca^2+^ imaging has therefore become a widely used approach for probing dendritic activity^9–12^. Yet, voltage and Ca^2+^ carry distinct information: membrane potential reflects many ionic and synaptic currents, while intracellular Ca^2+^ is additionally shaped by strong buffering, uptake by mitochondria^13^, and amplification through Ca^2+^-induced Ca^2+^ release from endoplasmic reticulum^14^. Similar Ca^2+^ waveforms can arise from distinct electrical patterns (or with no concurrent electrical event at all^15,16^), and subthreshold voltage dynamics often have no detectable Ca^2+^ signal. Thus, the relation between dendritic voltage and Ca^2+^ may be time dependent, history dependent, stochastic, and/or influenced by other dynamical variables.

Slice physiology established several important features of dendritic voltage-Ca^2+^ coupling. In CA1 pyramidal neurons, the extent to which somatic Na⁺ spikes backpropagate into the apical dendrites governs the spatial extent of dendritic Ca^2+^ entry, and this coupling depends on firing history^2,17–19^. Other slice experiments showed that clustered synaptic input can evoke Ca^2+^ excitations confined to apical dendrites^20^. During dendritic NMDA spikes and plateau potentials, Ca^2+^ currents drive large, long-lasting depolarizations ^2,18,20^ and burst firing at the soma^21^. *In vivo*, NMDA-dependent plateau potentials are associated with burst firing^22–24^ and plasticity^25–28^.

Given the different dynamical regimes of neural activity in slices vs. *in vivo*, one may not confidently extrapolate the voltage–Ca^2+^ mapping from reduced preparations. Only a few studies have directly measured dendritic voltage and Ca^2+^ simultaneously *in vivo*^29–31^, and these studies only probed small dendritic regions or discrete sites (**Supplementary Table 1**). To understand the relation between spike back-propagation, local inputs, and dendritic excitations, one would like to map simultaneous voltage and Ca^2+^ across the dendritic tree.

Hippocampal CA2 pyramidal neurons provide a useful system for addressing these questions. Though the electrophysiology of CA2 pyramidal cells is not as thoroughly studied as those in neighboring CA1 and CA3, dendrites of CA2 pyramidal cells are known to be excitable^32,33^, and distal apical excitations are implicated in plasticity of inputs from entorhinal cortex^34^. The apical dendrites of CA2 pyramidal cells branch close to the soma^32,33^, increasing the possibility of branch- specific electrical signaling and Ca^2+^ dynamics. CA2 neurons also show place field firing during spatial navigation^35^, providing a behavioral setting in which dendrite recruitment can be related to spatial coding.

Here, we developed Patterned Illumination Spinning disk COnfocal microscopy (PISCO) to simultaneously map membrane voltage and Ca^2+^ dynamics across nearly the entire dendritic arbor of individual CA2 pyramidal neurons in navigating mice. We used these measurements to determine the spatial footprints of voltage and Ca^2+^ excitations, to quantify the relation between local voltage waveforms and Ca^2+^ amplitude across compartments and behavioral states, and to examine how global spike back-propagation interacts with branch-specific depolarization during regenerative dendritic excitations. We identified a nonlinear and position-dependent, but deterministic, voltage-Ca^2+^ relationship. Our results support a model in which conjunction of back-propagating Na^+^ spikes and local dendritic input together recruit branch-specific Ca^2+^ electrogenesis that contributes to dendritic plateau potentials and somatic bursting.

## Results

### Simultaneous voltage and Ca^2+^ imaging in dendrites of behaving mice

The PISCO system combined patterned illumination with spinning disk confocal microscopy to provide kilohertz frame-rate optically sectioned imaging in two colors (**Fig. 1a**, **Fig. S1**). Separate digital micromirror devices (DMDs) patterned the red (607 nm, for voltage imaging) and blue (488 nm, for Ca^2+^ imaging) excitation lasers. These beams were then combined and re-imaged onto the microlens array of a customized spinning disk confocal microscope (**Methods**). The DMD patterns were designed to restrict illumination to the dendrites of interest, and to block illumination of interstitial regions.

**Fig. 1.**
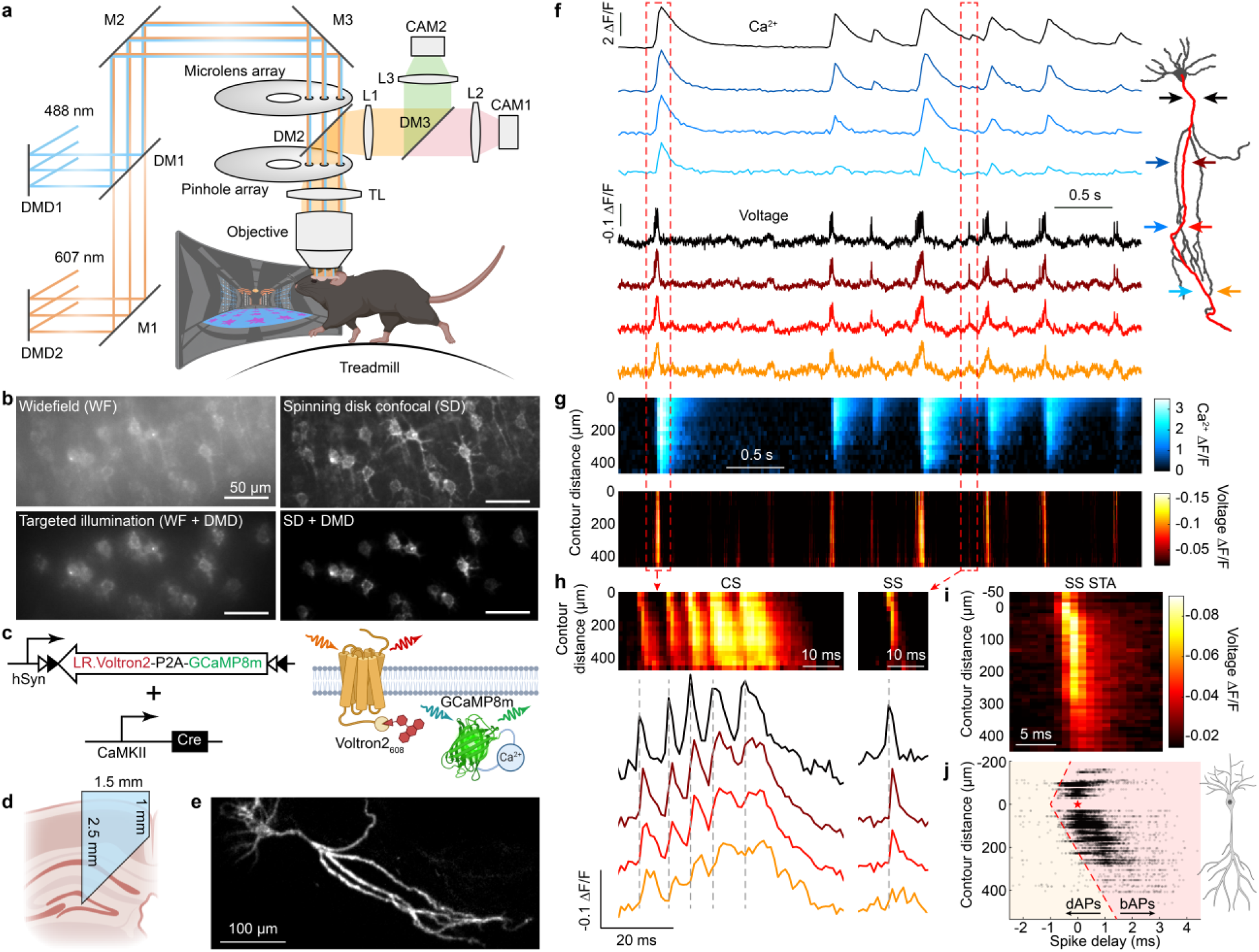
Simultaneous voltage and Ca^2+^ imaging across dendritic arbors of mice walking in VR. **a**, The PISCO system combines micromirror-based patterned illumination with spinning disk confocal microscopy for high-speed dual-wavelength imaging in tissue. DMD: digital micromirror device; M: mirror; DM: dichroic mirror; L: lens; TL: tube lens; CAM: camera. Relay lenses omitted for simplicity; full diagram in **Fig. S1**. **b,** Images of soma-targeted Voltron2-JF608 expression under different optical configurations. **c**, Genetic construct for sparse co-expression of the voltage indicator Voltron2 and Ca^2+^ indicator jGCaMP8m. **d**, An implanted microprism provided optical access to the hippocampus. **e**, Single-plane Voltron2-JF608 image of a hippocampal pyramidal neuron acquired *in vivo*. **f**, Simultaneously recorded Ca^2+^ (top) and voltage (bottom) dynamics from soma to distal dendrite (path highlighted on the right) *in vivo*. **g**, Kymographs of Ca^2+^ (top) and voltage (bottom) dynamics along the same dendritic branch in **f**. **h,** Zoomed-in voltage kymographs (top) and traces (bottom) of a complex spike (CS, left) and a simple spike (SS, right) along the same dendritic branch shown in **f**. **i,** Spike-triggered-average (STA) voltage kymograph of SSs from the cell shown in **f** (*n* = 56 spikes). **j,** Time delay of dendrite- detected spikes relative to the corresponding somatic spike (*n* = 16,111 dendrite-soma spike pairs from 10 cells, 7 mice). Spike timing with sub-frame resolution was estimated using spline interpolation (**Methods**). The red dashed lines indicate the classification boundary between bAPs and putative dAPs. Dendritic spikes occurring earlier than this boundary (left) were classified as putative dAPs, whereas those occurring later (right) were classified as bAPs. Negative contour distance indicates basal dendrite.

The spinning disk confocal and the DMD-patterned illumination worked synergistically: the spinning disk pinholes rejected light from outside the focal plane; the DMD patterning reduced diffuse background from multiply-scattered photons. Together, these two approaches achieved significantly higher contrast and signal-to-noise ratio compared to patterned illumination or spinning disk alone (**Fig. 1b**, **Fig. S1h**). Similar levels of contrast enhancement have been demonstrated with other hybrid confocal/targeted illumination systems^36^.

In the image path, a dichroic beamsplitter directed the Ca^2+^ signal to one camera and the voltage signal to a second camera. To achieve imaging at 1 kHz, we implemented phase-locked hardware synchronization between the voltage-imaging camera frame-clock and the spinning disk rotation^37^. We also implemented post-acquisition computational correction for slight variations between the twelve spinning disk sectors (**Fig. S2**, **Methods**). The Luminos software provided sub-pixel mapping of DMD to camera coordinates and microsecond-precision coordination of the whole experiment^38^.

The PISCO system achieved an optically sectioned imaging throughput of ∼2×10^8^ voxels/s, more than 50-fold higher than previous random-access two-photon methods used for dendritic imaging (**Supplementary Table 1**)^29–31^. This much-improved throughput was essential for simultaneous functional measurements across the dendritic arbor of hippocampal pyramidal neurons *in vivo*.

We co-expressed the red chemogenetic voltage indicator Voltron2-JF608 and the green Ca^2+^ indicator jGCaMP8m in hippocampal pyramidal cells of adult mice (**Fig. 1c**, **Methods**)^39,40^. A Lucy-Rho sequence was added to Voltron2 to improve its trafficking in dendrites^41,42^. We implanted a custom microprism to provide optical access to a sagittal transect of the hippocampus (**Fig. 1d**, **Methods**)^43^. While the prism field of view spanned CA1 and CA2, neurons suitable for dendrite-wide dual-color imaging were predominantly located in CA2. Dendritic trees of individual early-bifurcating CA2 pyramidal cells (**Fig. S3**)^32,44^ were clearly resolved from soma to apical tuft (**Fig. 1e**).

We verified via immunostaining for reactive astrocytes that the tissue damage was confined to ∼ 30 µm adjacent to the prism face (**Fig. S4**), consistent with prior microprism studies demonstrating stable chronic optical access and preserved tissue health beyond the interface zone^45,46^. We imaged neurons at distances of 50 – 150 μm from the prism face, where reactivity was minimal. The prism implantation location and orientation were selected to minimize interference with axonal connections to the recorded neurons^47^. Recent work using this prism preparation found no detectable differences in neuronal firing or place-field (PF) properties relative to the dorsal cannula preparation commonly used for *in vivo* hippocampus imaging^47^.

We combined the optical and molecular tools to map the voltage (1 kHz) and Ca^2+^ (41.5 Hz) dynamics across the dendritic trees of pyramidal neurons. During imaging, mice walked along a linear track in a novel 2-m virtual reality (VR) visual environment with a water reward once per lap (**Methods**). Our data clearly resolved subthreshold voltage fluctuations, spikes, and the accompanying Ca^2+^ dynamics (**Fig. 1f, g**, **Supplementary Video 1**). Recordings typically lasted 10 minutes (**Fig. S5**). During this time, photobleaching caused the Voltron2 fluorescence to decay to 71 ± 5% of its initial amplitude (mean ± s.d., *n* = 13 cells), and the jGCaMP8m fluorescence to decrease to 75 ± 6% of its initial amplitude. These decays were corrected in post-processing (**Methods**). Though Ca^2+^ events were detected based on the jGCaMP8m signal alone, all detected Ca^2+^ events (*n* = 1832 events over 13 cells, 10 mice) had a corresponding electrical excitation, confirming that false-positive detections were unlikely. In the apical tuft, where jGCaMP8m fluorescence was dimmest, complex spike-associated Ca^2+^ events had a signal-to-noise ratio (SNR) of 17.7 ± 4.6 (mean ± s.d.), 5.9-fold larger than the event-detection threshold set at SNR = 3 (**Methods**). The full image processing pipeline is described in **Methods** and **Fig. S6**.

We classified somatic excitations by their spike timing and subthreshold voltage dynamics. Single action potentials separated from neighboring events by > 25 ms were classified as simple spikes (SSs, **Fig. 1h,i**). Groups of spikes in which all inter-spike intervals were ≤ 25 ms were classified as bursts. Bursts with 4 or more spikes riding atop a specified subthreshold depolarization at the soma were classified as complex spikes (CSs, **Fig. 1h**, **Methods**).

We next used the kilohertz voltage recordings to determine the propagation direction of simple spikes. Voltage spikes were detected independently in each subcellular compartment and linked across neighboring compartments when their peak times differed by ≤ 1 ms (**Methods**). Of Na^+^ spikes detected in soma and dendrites, most first appeared near the soma and propagated into apical and basal compartments with distance-dependent delays (98.6%; 15,880 out of 16,111 events, 10 cells, 7 mice; **Fig. 1h-j**). We classified these as back-propagating action potentials (bAPs) and calculated a spike propagation speed of 0.22 ± 0.01 m/s for apical dendrites and 0.27 ± 0.06 m/s for basal dendrites (**Fig. S7**). The remaining events (1.4%) peaked in a dendrite earlier than expected from these propagation speeds. These rare events were classified as putative dendrite- initiated spikes (dAPs, **Fig. 1j**).

## Distal apical Ca^2+^ is primarily associated with complex spikes

We then explored the Ca^2+^ footprints of different types of voltage events. Consistent with previous studies^2,17^, SSs reliably induced Ca^2+^ transients at the soma and proximal dendrites (**Fig. 2a, b**). The peak voltage from an SS (measured as ΔF/F) decayed with distance along the apical dendrites with a length constant of 303 ± 49 μm (mean ± s.d., *n* = 900 events, 11 neurons, 8 mice, **Fig. S7**). The peak Ca^2+^ signal of the same events decayed with a length constant of 184 ± 36 μm (**Fig. S7**). While the voltage signal from SS events was clearly detectable in the most distal dendrites (483 μm from the soma), there was no detectable Ca^2+^ signal in these dendrites. In contrast, CSs engaged larger amplitude and more spatially extended Ca^2+^ signals, including in distal apical dendrites. The ratio of mean CS Ca^2+^ amplitude to mean SS Ca^2+^ amplitude was an increasing function of distance from the soma (perisoma: 4.3 ± 0.3; distal dendrite: 10.7 ± 2.6, mean ± s.e.m., *n* = 11 neurons, 8 mice. **Fig. 2b**).

**Fig. 2.**
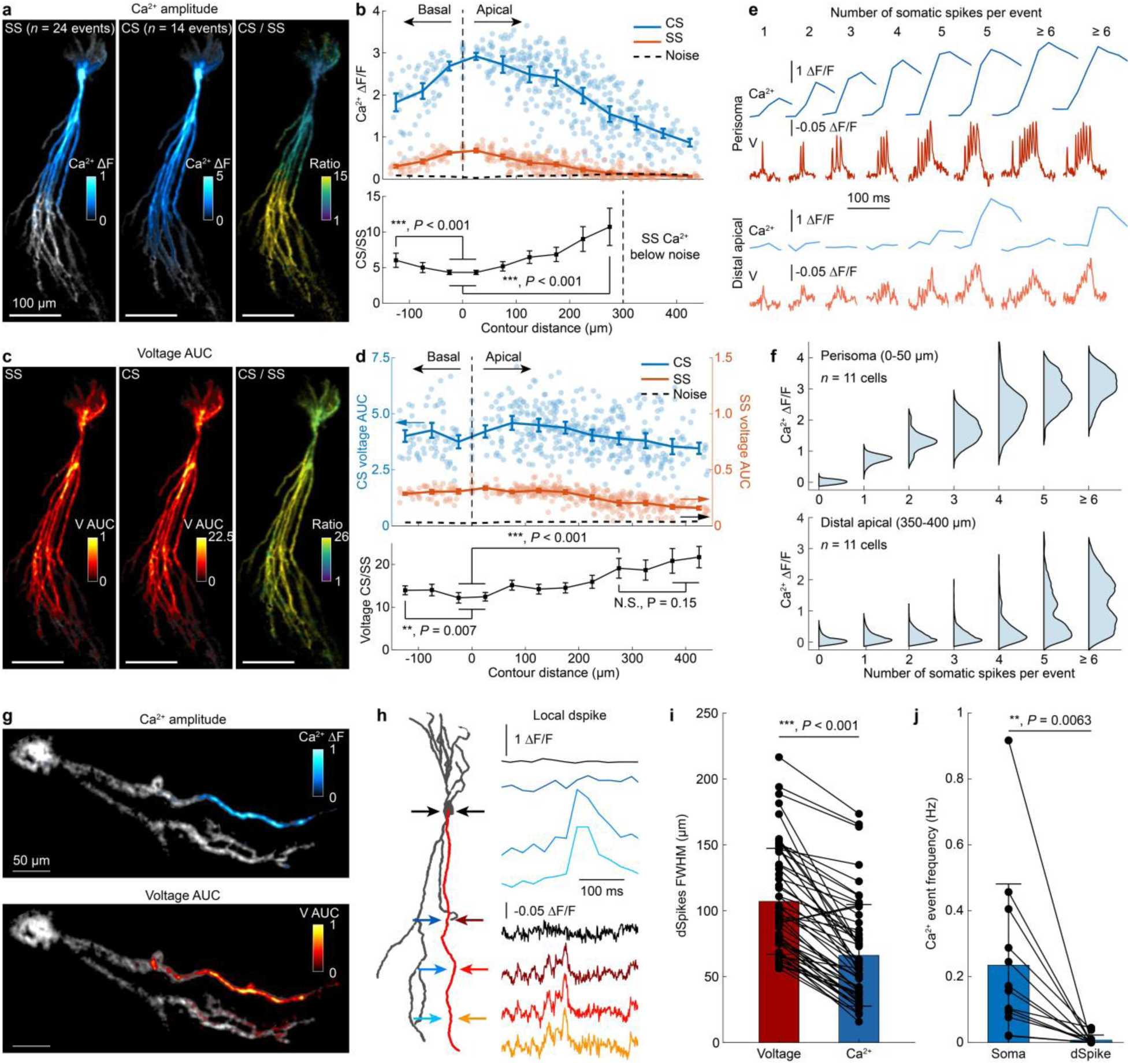
Distal apical Ca^2+^ is primarily associated with complex spikes. a, Average Ca^2+^ spatial profiles for SS (left) and CS (middle) events, overlaid on a structural image of the neuron. The right panel shows the ratio between CS and SS Ca^2+^ amplitudes. **b,** Top: Ca^2+^ ΔF/F for CS and SS events as a function of contour distance from the soma. The noise floor was determined by 3×s.d. of the silent period (i.e. no Ca^2+^ spikes). Bottom: Ratio of mean CS to SS Ca^2+^ ΔF/F as a function of distance from the soma. Data are presented as mean ± s.e.m. One-way analysis of variance (ANOVA). **c,** Average voltage AUC spatial profiles for SS (left) and CS (middle) events, overlaid on a structural image of the neuron. The right panel shows the ratio between CS and SS voltage AUCs. **d,** Top: Voltage AUC for CS and SS events as a function of contour distance from the soma. The noise floor was determined by 3×s.d. of the silent period (i.e. no voltage spikes). Each point in the upper panels of **b** and **d** represents the average value across all events of the indicated type in a subcellular compartment (*n* = 11 cells, 8 mice). Bottom: Ratio of mean CS to SS voltage AUC as a function of distance from the soma. Data are presented as mean ± s.e.m. One-way analysis of variance (ANOVA). N.S., not significant. **e,** Representative near-soma (top) and distal dendrite (bottom) Ca^2+^ (blue) and voltage (red) dynamics for events with different numbers of spikes. Events with ≥ 5 spikes with and without an associated apical Ca^2+^ signal are shown separately. **f,** Distribution of near-soma (top) and distal dendrite (bottom) Ca^2+^ ΔF/F for events with different numbers of spikes (*n* = 11 cells, 8 mice, 239 – 1058 events per category). **g,** Maps of Ca^2+^ ΔF/F (top) and voltage AUC (bottom) for a representative local dSpike, overlaid on a grayscale structural image of the neuron. **h,** Representative Ca^2+^ and voltage dynamics during the local dSpike in **g**. **i**, Spatial extent (full width at half-maximum, FWHM) of local voltage and Ca^2+^ dSpikes. Data are presented as mean ± s.d.. Each point represents one local dSpike event. (*n* = 54 events, 4 cells, 4 mice). Paired t-test. The voltage dSpikes were 52 ± 17 ms in duration (mean ± s.d.). **j,** Frequency of somatic Ca^2+^ events and local Ca^2+^ dSpikes. Data are presented as mean ± s.d. and each point represents one cell (*n* = 13 cells, 10 mice). Paired t-test.

We compared the Ca^2+^ spatial profiles to the corresponding voltage profiles. We quantified voltage event size via area-under-the-curve (AUC) of ΔF/F vs. t (**Methods**). Despite a >10-fold difference in voltage AUC between SS vs. CS events, the shapes of the AUC spatial profiles were similar: both event types extended to the most distal dendrites in the field-of-view (up to 483 µm contour distance) and varied smoothly with distance from the soma (**Fig. 2c, d**). The dimensionless ratio of CS to SS voltage AUC showed a modest increase with distance from the soma (perisoma: 12.3 ± 1.1; distal dendrite: 19.1 ± 2.3, mean ± s.e.m., *n* = 11 neurons, 8 mice). Thus the modest differences in CS/SS voltage spatial profiles corresponded to strong distance dependence of CS/SS Ca^2+^ signals.

We next extended the analysis across all electrical events with different number of spikes. Near the soma, Ca^2+^ amplitude was an increasing, but sub-linear, function of the number of spikes in an electrical event (**Fig. 2e, f**), consistent with recent simultaneous patch-clamp and jGCaMP8m recordings *in vivo*^40^. In distal apical dendrites (> 350 μm from the soma), events with ≤ 3 spikes were not accompanied by detectable Ca^2+^ signal, and almost all detectable Ca^2+^ accompanied CS events (**Fig. 2a, b, e, f**). However, a CS did not invariably coincide with Ca^2+^ recruitment in a given distal branch, resulting in a clear bimodal distribution of Ca^2+^ ΔF/F amplitude (apical Ca^2+^ transients in 51 ± 22% of CSs, mean ± s.d., *n* = 550 events, 29 branches, 11 neurons, **Fig. 2e, f**, Fig. **S8**). This bimodal recruitment is consistent with a regenerative process in distal dendrites (e.g. Ca^2+^ electrogenesis coupled with voltage-dependent calcium conductance), as proposed in classic slice physiology experiments.^48,49^

## Isolated dendritic voltage and Ca^2+^ events without concurrent somatic events are rare

We also observed rare voltage and Ca^2+^ events co-localized to individual dendritic branches (dSpikes), without detectable somatic voltage or somatic Ca^2+^ signals (*n* = 57 events in 2 recording hours, 4 out of 13 cells, 4 out of 10 mice, **Fig. 2g, h**). The local voltage dSpikes had a wider spatial footprint than the corresponding Ca^2+^ dSpikes (full width at half-maximum for voltage: 107 ± 40 µm, Ca^2+^: 66 ± 39 µm, mean ± s.d., **Fig. 2i**). The rate of Ca^2+^ dSpikes was 29-fold lower than that of the somatic Ca^2+^ events (0.008 ± 0.016 Hz vs. 0.23 ± 0.25 Hz, mean ± s.d., *n* = 13 cells, **Fig. 2j**). Our observation that ∼97% of dendritic Ca^2+^ events co-occurred with somatic Ca^2+^ events (but not *vice versa*) is consistent with prior reports in CA1^26^, CA3^50^, and L5 pyramidal neurons^11,51–53^ *in vivo*.

In summary, whereas voltage events appeared largely delocalized over the dendritic tree, Ca^2+^ signals followed a hierarchical structure: Ca^2+^ events were almost always largest at the soma, with footprints that extended to variable distances along the dendritic tree.

## Broadly shared dendritic voltage coexists with branch-specific Ca^2+^

The variability in distal apical Ca^2+^ across CS events motivated us to examine the spatial organization of voltage and Ca^2+^ during individual events. Distal apical branches with broadly similar CS voltage signals sometimes had substantially different Ca^2+^ amplitudes (**Fig. 3a**). **Fig. 3b** illustrates two successive CS events, labeled α and β, which showed opposite Ca^2+^ success and failure patterns in two dendrites, D1 and D2. For each event, the accompanying voltage signal was only slightly larger in the Ca^2+^-positive branch than in the Ca^2+^-negative branch. This pattern of branch-specific variations in Ca^2+^ and broadly shared voltage re-occurred across all CS events in the recording (**Fig. 3a-f**, **Supplementary Videos 2, 3**). The Ca^2+^ failures often occurred near branch points, though the voltage signals decayed smoothly across the branch points (**Fig. 3c**, **g**, **Fig****. S9**). Similar branch-specific recruitment of dendritic Ca^2+^ signals has been observed in both CA1 and cortical pyramidal neurons in brain slices^2,19,54^ and *in vivo*^24,27,51^. Our simultaneous arbor- wide measurements reveal that this branch specificity is substantially stronger for Ca^2+^ than for voltage during the same events *in vivo*.

**Fig. 3.**
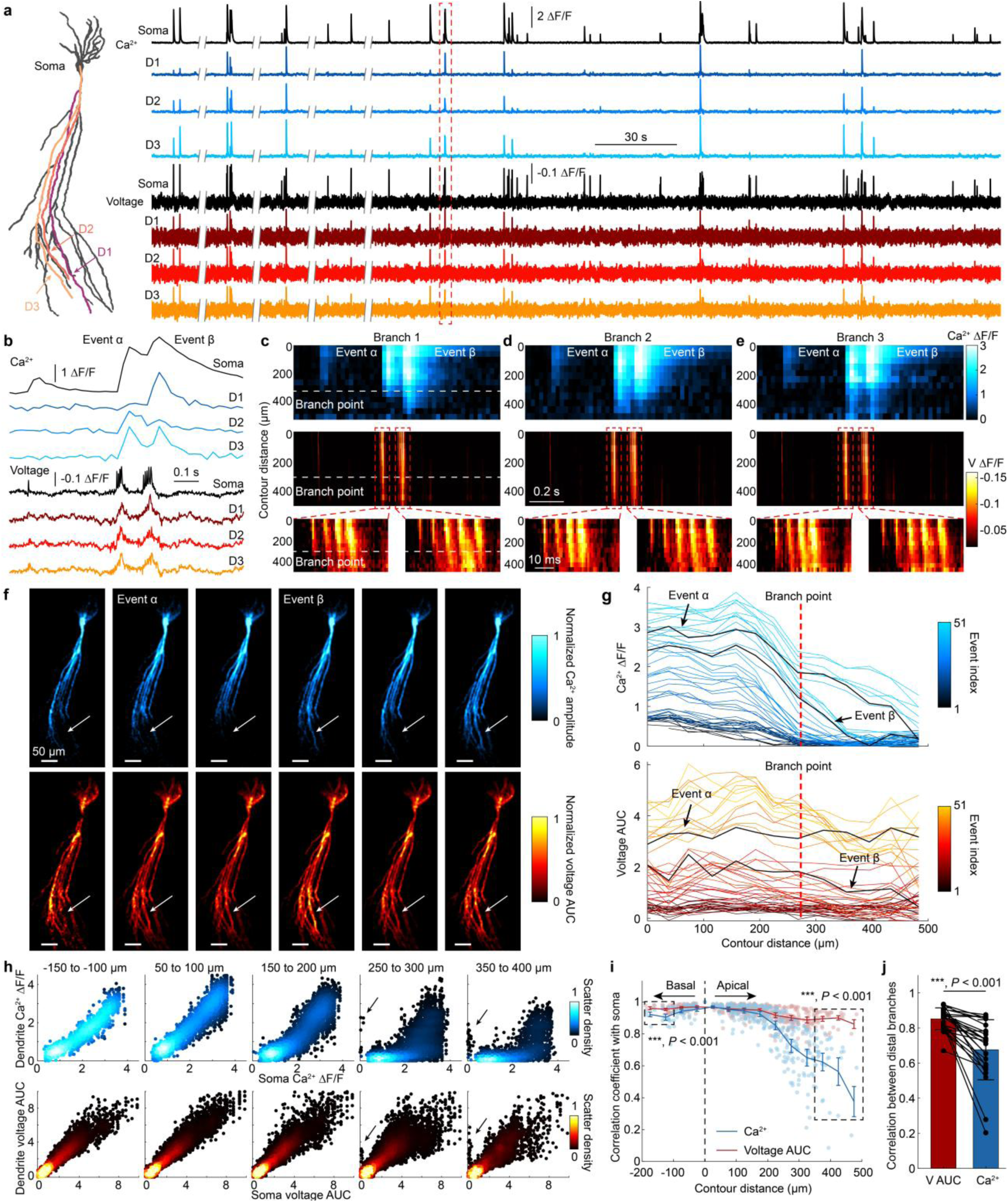
Broadly shared dendritic voltage coexists with branch-specific Ca^2+^. a, Representative Ca^2+^ (top) and voltage (bottom) dynamics in the soma and three distal dendrites (highlighted on the left). **b,** Zoomed-in Ca^2+^ and voltage dynamics in the red box in **a**. Events α and β exhibit branch-specific Ca^2+^ signals. **c-e** Kymographs of Ca^2+^ (top) and voltage (middle) for events α and β along the same 3 branches as in **a**. The bottom panels show the zoomed-in voltage kymographs for the two events. **f,** Maps of normalized Ca^2+^ ΔF/F (left) and normalized voltage AUC (right) for six representative events. For each event, Ca^2+^ and voltage maps were separately normalized to their corresponding somatic values. White arrows indicate D1, which shows distinctive Ca^2+^ signals between events. **g,** Spatial profiles of Ca^2+^ ΔF/F (top) and voltage AUC (bottom) along D1. Each line represents one event. Event coloring is sorted based on Ca^2+^ amplitude in the soma. **h,** Dendritic vs. somatic Ca^2+^ ΔF/F (top) and voltage AUC (bottom) at different contour distances. The black arrows indicate local dSpikes. Each data point represents one event in a single subcellular compartment (*n* = 11 cells, 8 mice). **i,** Correlation coefficients of dendritic Ca^2+^ with somatic Ca^2+^, and dendritic voltage AUC with somatic voltage AUC, as a function of contour distance. Data are presented as mean ± s.e.m. and each point represents one subcellular compartment (*n* = 11 cells, 8 mice). Paired t-test. **j**, Correlation coefficients of CS Ca^2+^ and voltage AUC between pairs of distal branches within each cell. Data are presented as mean ± s.d. and each point represents one pair of distal branches within the same cell (11 – 159 CS events for each pair of distal branches, 25 pairs of distal branches, 11 cells). Paired t-test.

To quantify this effect, we compared event-by-event voltage AUC and Ca^2+^ ΔF/F amplitude at soma vs. dendrites. Across events with different numbers of spikes and different subthreshold amplitudes (including all SS and CS events), the dendritic and somatic voltage AUC were highly correlated across all basal and apical branches, with the correlation decreasing only modestly with soma-dendrite contour distance (**Fig. 3h, i**, **Fig****. S10**). In contrast, the correlation between dendritic and somatic Ca^2+^ ΔF/F decreased sharply in distal apical dendrites beyond ∼200 µm from the soma (**Fig. 3h, i**, **Fig****. S10**). Similarly, the voltage AUC between pairs of distal apical dendrites from the same cell was also highly correlated, while the pairwise correlation between the corresponding Ca^2+^ ΔF/F amplitudes was significantly lower (**Fig. 3j**). In summary, voltage signals were largely shared across the dendritic arbor, whereas Ca^2+^ footprints showed stronger branch-to-branch and event-to-event heterogeneity. This difference in spatial structures raised the question of to what extent the Ca^2+^ and voltage signals had a fixed relationship vs. varied independently due to noise or un-measured variables.

## A simple biophysics-inspired model predicts local Ca^2+^ from local voltage *in vivo*

Although local voltage and Ca^2+^ are bidirectionally coupled (via voltage-gated Ca^2+^ currents and Ca^2+^ electrogenesis), nonetheless one might (or might not) find a predictive empirical relation between these quantities. We plotted local Ca^2+^ ΔF/F vs. local voltage AUC across all excitation events (including SS, CS, and dSpikes) and all measured sub-cellular compartments. The voltage- Ca^2+^ relations transitioned from sublinear at and near the soma to sigmoidal with an apparent threshold in distal apical dendrites (**Fig. 4a**, **Fig. S11**). Prior efforts to model the voltage-Ca^2+^ relation aspired to multi-component realistic descriptions^55,56^, but the visually apparent trends in our data suggested that a simple semi-empirical fit might suffice.

**Fig. 4.**
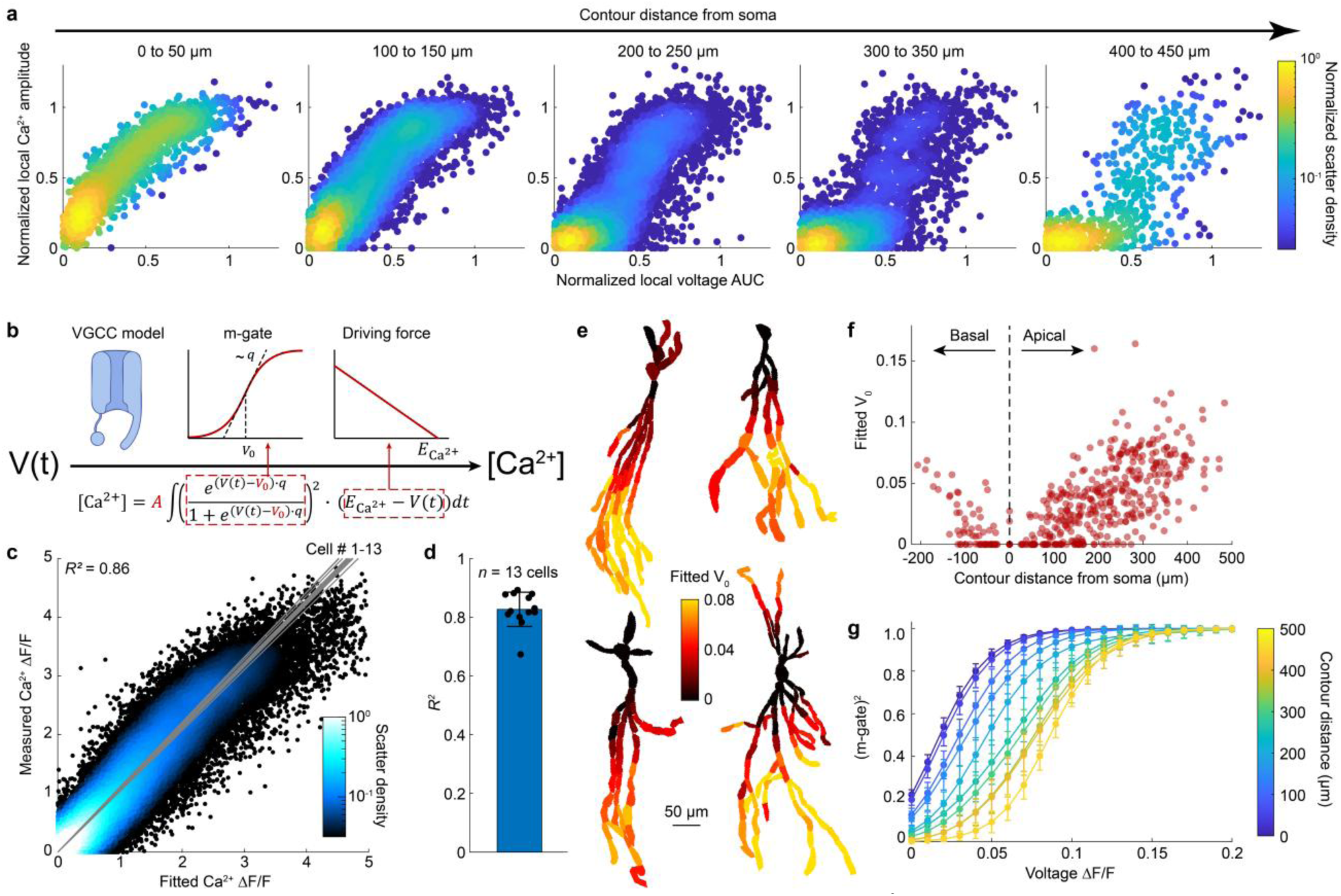
A simple biophysics-inspired model predicts local Ca^2+^ from local voltage *in vivo*. a, Normalized. Ca^2+^ amplitude vs. normalized local voltage AUC at different contour distances from the soma. Each datapoint represents one event in a single subcellular compartment (*n* = 11 cells, 8 mice). **b,** Scheme showing the biophysics-inspired model that predicts Ca^2+^ ΔF/F from voltage. **c,** Measured Ca^2+^ ΔF/F vs. model-predicted Ca^2+^ ΔF/F from local voltage waveform. Each point represents one event in a single subcellular compartment (*n* = 13 cells, 528 compartments, 1775 spiking events, 53,592 measurements). Gray lines: linear fit for each cell. **d,** Statistics of the *R*^2^ values for all cells in **c**. Data are presented as mean ± s.d. and each point represents one cell. **e,** Representative spatial structures of fitted *V*_0_ values in four cells. **f,** Scatterplot of fitted *V*_0_ values as a function of contour distance from soma. Each point represents one subcellular compartment (*n* = 11 cells). **g,** Model-derived (m-gate)^2^ as a function of voltage ΔF/F for apical dendrites grouped by contour distances. Data are presented as mean ± s.e.m.

We hypothesized that the voltage-Ca^2+^ relation might be captured by a family of sigmoidal curves resembling ion channel activation gates. We made a minimal model inspired by voltage-dependent gating of a hypothetical voltage-gated Ca^2+^ channel (VGCC, **Fig. 4b**, **Methods**)^57,58^, though we emphasize that this model is not molecularly specific: it seeks a quantitative description of the voltage-Ca^2+^ relationship, capturing contributions from VGCCs, NMDA receptors, intracellular Ca^2+^ handling, and Ca^2+^ electrogenesis, with a small number of parameters. We described the channel gating by a sigmoidal function *m*_∞_(*V*), and assumed that the channel open probability followed *P_open_* = *m*^2^ .^54,59^ This model assumed rapid channel activation and ignored channel inactivation (i.e. no *h* gate). We assumed a Ca^2+^ influx rate *i*_Ca2+_ ∝ *P_open_* × (*E*_Ca2+_ − *V*(*t*)), where (*E*_Ca2+_ − *V*(*t*)) is the driving force for Ca^2+^ influx. Since the voltage events were much faster than the Ca^2+^ events, and the Ca^2+^ upstroke was much faster than the return to baseline, we treated the voltage events as an impulsive Ca^2+^ influx and we did not model Ca^2+^ efflux, i.e. the model converted the function *V*(*t*) for each event into a prediction of the peak Ca^2+^ ΔF/F.

We estimated the effective driving-force, *E*_Ca2+_, (in units of ΔF/F) and the gating steepness *q* (units (ΔF/F)^-1^) based on published biophysical measurements and the Voltron2 F vs. V relation (**Methods**). We held these parameters fixed across all cells and compartments. We then fitted two parameters for each subcellular compartment: the activation threshold *V*_0_ (units of ΔF/F) and an overall scaling factor, *A*. We performed the fit for *n* = 13 neurons, with 13 – 79 compartments per cell and 12 – 550 events per cell (1775 total events, 528 total compartments, 53,592 total predictions). This compact model successfully predicted Ca^2+^ event amplitudes across all compartments in all cells with *R*^2^ = 0.86 (**Fig. 4c, d**, **Fig****. S12**). Saturation of the jGCaMP8m signal at high [Ca^2+^] might account for the model’s slight overestimate of Ca^2+^ ΔF/F amplitude for the largest events. Although local dSpikes were excluded from parameter fitting, the model also predicted dSpike Ca^2+^ amplitudes from the accompanying local voltage waveforms without any adjustments of parameters (**Fig. S13**, *R^2^* = 0.69). Our model established a 1:1 deterministic relation between local voltage dynamics and peak local Ca^2+^ in each dendritic segment.

Though the activation threshold *V*_0_ was fit independently for each dendritic compartment, this parameter showed regular trends across cells. *V*_0_ increased with distance from the soma in both apical and basal dendrites (**Fig. 4e-g**). This trend captured the transition from sub-linear V-Ca^2+^ coupling near the soma to sigmoidal and increasingly step-like relations in the apical dendrites. The change in shape of the voltage-Ca^2+^ relationships could not be trivially explained by position- dependent changes in sensitivity of either the voltage or Ca^2+^ recordings. Rather, this finding implies a gradient toward increasing threshold for activation of Ca^2+^ influx with increasing distance from the soma. In distal apical dendrites, the steep voltage-Ca^2+^ nonlinearity was sufficient to account for the co-existence of broadly shared voltage signals and branch-specific Ca^2+^ signals during the same events. We emphasize that this V-Ca^2+^ relationship is an empirical mapping between jointly measured signals rather than a unidirectional causal transfer function.

## Pre-spike apical depolarization correlates with enhanced apical CS Ca^2+^

We next asked what caused the variations in apical CS voltage waveforms and the associated Ca^2+^ success or failure? Based on prior studies in acute slices,^57,60,61^ we hypothesized that pre-spike excitatory inputs could affect the reserve of glutamate-bound NMDA receptors. A sufficient pool of ‘primed’ NMDA receptors could support voltage-gated Mg^2+^ unblock and regenerative Ca^2+^ influx.

In each distal branch and for each CS event, we quantified local depolarization leading up to the first CS spike, by calculating the voltage AUC during the 100 ms preceding the event (pre-AUC). High-Ca^2+^ events showed, on average, a higher pre-AUC compared to low-Ca^2+^ events (high Ca^2+^: pre-AUC = 0.44 ± 0.54; low Ca^2+^: pre-AUC = -0.09 ± 0.50, *P* = 2.6 × 10^-6^, *n* = 657 events, 20 distal branches, 8 neurons, 7 animals, **Fig. 5a, b**, **Fig****. S14**, **Methods**). Thus, during CS events small apical depolarizations lead to large local Ca^2+^ signals, as one would want for an associative plasticity mechanism. A similar effect was reported in Purkinje cell dendrites^31^.

**Fig. 5.**
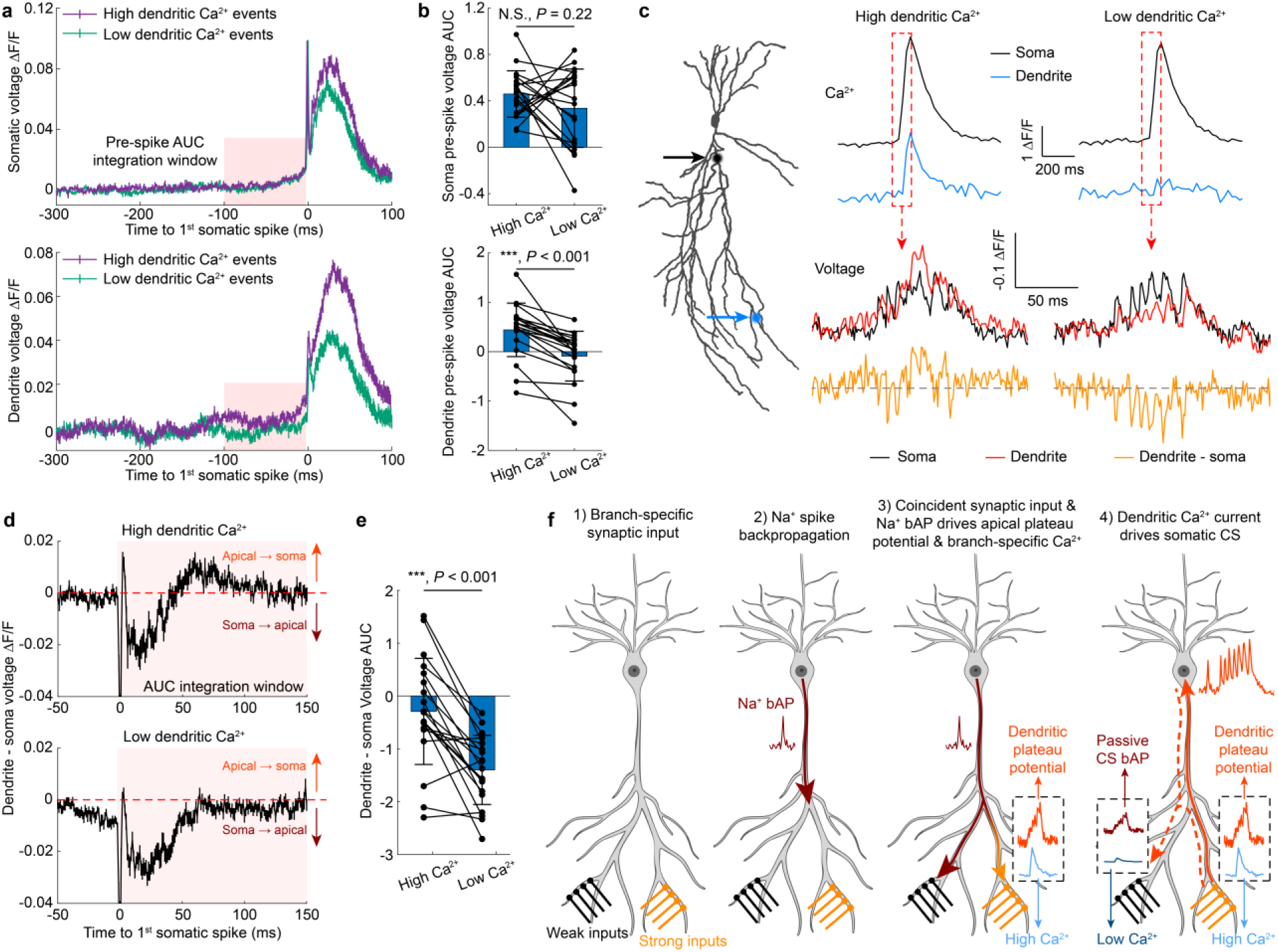
Conjunctive apical depolarization and somatic spiking drives apical plateaus, Ca^2+^ influx, and CS events. a, Average somatic (top) and dendritic (bottom) voltage waveforms for CS events with high (purple) vs. low (green) apical dendritic Ca^2+^. Data are presented as mean ± s.e.m. **b,** Pre-spike voltage AUC from -100 to -2 ms in the soma (top) and dendrite (bottom) for CS events with high vs. low dendritic Ca^2+^. Data are presented as mean ± s.d. and each point represents the average value from a distal dendrite. Paired t-test. **c,** Representative somatic and distal apical Ca^2+^ (top) and voltage (bottom) traces for high (left) and low (right) dendritic Ca^2+^ events. The dashed gray line indicates zero. The orange trace reports the voltage gradient between dendrite and soma. **d,** Average *dendrite* - *soma* voltage gradients for dendritic events with high (top) and low (bottom) apical Ca^2+^, aligned to the first somatic spike. All events in the corresponding categories from each dendritic branch were first averaged, and the mean waveforms were then averaged across branches (See **Methods** for the definition of high and low Ca^2+^ events). Positive values indicate apical-to- soma current flow. Events were aligned to the peak of the first spike. Spikes after the first are not visible in the average waveform because the spike times varied between events. Data are presented as mean ± s.e.m. (657 events from 20 distal dendrites in 8 cells). **e,** Statistical analysis of *dendrite* - *soma* voltage AUC for dendritic events with high vs. low apical Ca^2+^. The AUC integration window is highlighted in **d**. Data are presented as mean ± s.d. and each point represents the average value from a distal dendrite. Paired t-test. **f,** Model for branch-specific regenerative dendritic recruitment during CSs. Conjunction of branch-specific synaptic excitation and back-propagating action potentials triggers a dendritic Ca^2+^ spike and plateau potential, which drive a CS at the soma. Branches in which NMDA receptors have bound glutamate due to pre-CS synaptic inputs contribute disproportionally to the plateau Ca^2+^ current and have the largest Ca^2+^ elevations.

## High-Ca^2+^ apical plateau potentials drive current into the soma

We then analyzed the consequences of apical excitations for somatic output. We compared the somatic and apical voltage waveforms during CSs with high vs. low apical Ca^2+^ (**Fig. 5c-e, Methods**). We used the dendrite − soma voltage difference to infer the direction of current drive^41^: negative values indicate soma-drives-dendrites; positive indicate dendrites-drive-soma. Both high and low dendritic Ca^2+^ events showed soma-drives-dendrites for the first several somatic spikes (approximately 40 ms), consistent with prior measurements in acute slices^41^ and the view that multiple bAPs are a necessary trigger for CS events (**Fig. 5c, d, Fig. S15**). In high-Ca^2+^ events, the gradient then reversed, with the distal apical dendrites more depolarized than, and hence driving, the soma. Low-Ca^2+^ CS events retained a slight soma-to-dendrite voltage gradient throughout (**Fig. 5c-e**). Thus, during CS events with high apical Ca^2+^, the apical dendrites switched from a sink to a source of somatic current.

Together, our data support a simple biophysical picture of a complex spike (**Fig. 5f**). Apical glutamate inputs prime NMDA receptors; conjunctive arrival of back-propagating action potentials removes the NMDAR Mg^2+^ block, triggering a regenerative Ca^2+^ influx. The resulting apical plateau potential drives somatic CS firing. Other apical branches with weaker pre-spike synaptic excitation experience passive CS back-propagation and have low Ca^2+^ signals. Our data provide *in vivo* validation of a model of bAP-activated Ca^2+^ electrogenesis previously proposed based on acute slice experiments^57,60,61^.

### Behavioral state and spatial context modulate event prevalence and distal apical recruitment

Finally, we asked how the dynamics described above manifest during behavior. Locomotion led to increased Ca^2+^ event amplitudes throughout the dendritic tree (**Fig. S16a**), but did not change the compartment-specific voltage-Ca^2+^ relation (**Fig. S16b**). During running we observed an elevated proportion of CS events, and the running-associated CS events had higher apical Ca^2+^ amplitude, compared to CS events during rest (**Fig. S16c, d**).

A subset of the recorded neurons showed place field (PF)-like firing patterns (5 place cells from 5 mice, of 13 measured cells). As with locomotion, PF context led to increased Ca^2+^ event amplitudes throughout the dendritic tree (**Fig. 6a**)^27^ relative to non-PF locations, but did not change the compartment-specific voltage-Ca^2+^ relation (**Fig. 6b**). PFs contained an elevated proportion of CS events, and the PF-associated CS events had higher apical Ca^2+^ amplitude compared to CS events outside the PF. Thus, both locomotion and PF activity were associated with a shift toward CS firing and toward enhanced apical Ca^2+^ during CSs, but did not change the underlying V-Ca^2+^ relation.

**Fig. 6.**
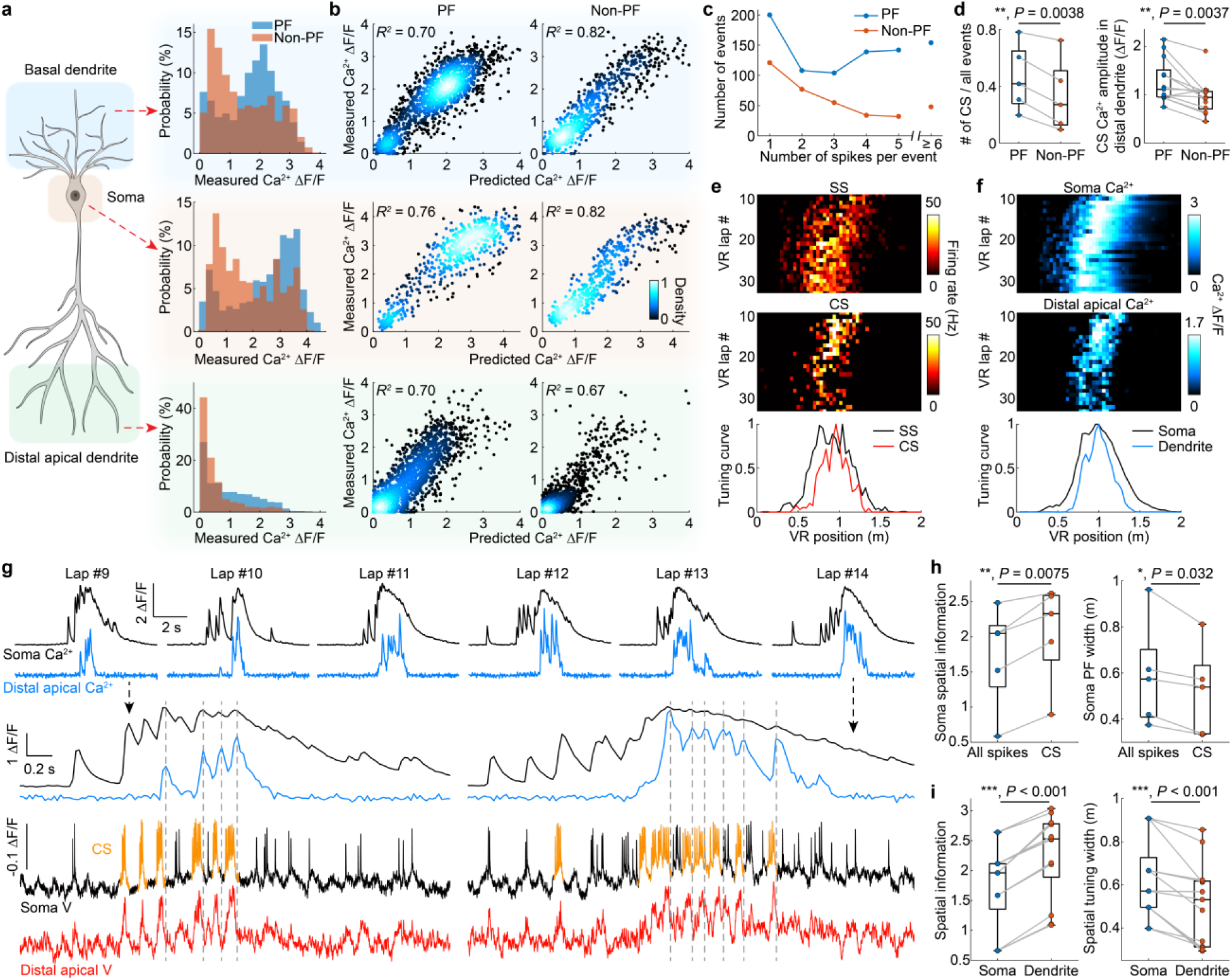
Brain state modulates event prevalence and structure but preserves V-Ca^2+^ mapping. a, Distribution of measured Ca^2+^ event amplitude within and outside the smatic place field (PF) for basal dendrites (top), soma (middle), and distal apical dendrites (bottom). *n* = 5 place cells. **b,** Measured Ca^2+^ ΔF/F vs. model-predicted Ca^2+^ ΔF/F for PF and non-PF events across different subcellular compartments. **c,** Number of events as a function of somatic spike count within and outside the somatic PF. **d,** Left: Fraction of events classified as CS. Each point represents one cell. Right: mean distal apical Ca^2+^ amplitude for PF and non-PF CS events. Each point represents one distal branch (*n* = 13 distal branches from 5 place cells). Paired t-test. **e,** Example somatic SS (top) and CS (middle) firing rate maps as a function of VR position and lap number. Bottom: spatial tuning curves of SS and CS. **f,** Example somatic (top) and distal apical (middle) Ca^2+^ ΔF/F maps for the same cell in **e**. Bottom: spatial tuning curves of soma and distal apical dendrite. **g,** Top: Example somatic (black) and distal apical (blue) Ca^2+^ traces across 6 VR laps. Bottom: Zoomed- in Ca^2+^ and voltage (soma: black, distal apical: red) traces for Lap #9 and Lap #14. Somatic CSs are highlighted in yellow. **h,** Somatic spatial information (left) and PF width (right) for all spikes vs. CSs only. Each point represents one cell. (*n* = 5 place cells from 5 mice). Paired t-test. **i,** Ca^2+^- based spatial information (left) and spatial tuning width of soma and distal apical dendrites, calculated from the corresponding Ca^2+^ traces (**Methods**). Each point represents one soma- dendrite pair. (*n* = 13 dendrite-soma pairs from 5 place cells). Paired t-test.

We next asked about compartment-specific spatial tuning. Somatic CS firing was more spatially selective than other spikes^47^, exhibiting higher spatial information and narrower tuning around the PF (**Fig. 6e, h**). Distal apical Ca^2+^ activity was also more spatially selective than somatic Ca^2+^ activity in place cells (**Fig. 6f, g**). Across distal apical branches, Ca^2+^-based spatial information was 1.4 ± 0.2 (mean ± s.d., *P* = 1.4 ×10^-7^) times the corresponding somatic value, and the spatial tuning width was 0.80 ± 0.15 (mean ± s.d., *P* = 4.6 ×10^-4^) times the somatic value (**Fig. 6i**). Similar differences between somatic and dendritic spatial Ca^2+^ tuning have been reported in CA1 place cells^26^. Our model predicts that CS events, and particularly high-apical-Ca^2+^ CS events, arise from conjunction of apical inputs and bAPs. We suggest that the conjunction of these two inputs is more tightly tuned than either input alone, leading to the sharper apical Ca^2+^ tuning observed here.

## Discussion

Dendrites must solve a fundamental problem: how to convert the rich but fleeting vocabulary of electrical signals into the slower biochemical language that governs synaptic plasticity. Conversion of electrical events into Ca^2+^ signals is the critical electrical-chemical coupling step. Here, by simultaneously mapping voltage and Ca^2+^ across the dendritic arbors of CA2 pyramidal neurons during spatial navigation, we provide what is, to our knowledge, the first dendrite-wide characterization of the voltage-Ca^2+^ coupling in a behaving mammal.

Dendritic voltage and Ca^2+^ had strikingly different spatial organization: voltage was broadly coordinated across the dendritic arbor, whereas Ca^2+^ showed pronounced branch-to-branch and event-to-event heterogeneity, particularly in distal apical dendrites. Nonetheless, the two signals had a deterministic and local 1:1 relationship (*R*^2^ ≈ 0.86), captured by a compact biophysics- inspired model. While the model predicts Ca^2+^ from voltage, the two variables are bidirectionally coupled: voltage activates Ca^2+^ conductances, and Ca^2+^ currents contribute to voltage. An increasingly steep voltage-to-Ca^2+^ transfer function in distal dendrites mapped subtle changes in voltage to large, nearly binary, changes in Ca^2+^, likely reflecting positive feedback via Ca^2+^ electrogenesis.

A surprising implication of our results is that, despite the biological complexity of Ca^2+^ handling, the presence of behavior-dependent modulation of event prevalence, and un-observed variables such as inhibitory and modulatory signals, Ca^2+^ and voltage appear to be deterministically coupled. Whether this deterministic relationship persists throughout long-term plasticity or across behavioral states beyond those we tested remains to be determined.

Dendrites have been modeled as independently excitable subunits capable of generating local Ca^2+^ spikes that may drive somatic activity^24,62–67^. Our findings in CA2 suggest a revised picture (**Fig. 5f**). Dendritic excitations rarely occurred on their own, and rarely triggered the first spike in a burst or CS. Rather, apical dendritic excitations arose from conjunction of bAPs and apical depolarization. These apical excitations then sourced current to the soma, driving somatic CS events.

Branch-specific excitatory synaptic input is a plausible source of the pre-spike dendritic depolarization. Glutamatergic input can provide local depolarization and prime NMDA receptors for subsequent Mg^2+^ unblock during a CS event. Slice experiments have shown that coincident bAP and synaptic inputs can amplify local voltage and Ca^2+^.^57^ However, voltage imaging alone does not identify the underlying conductances. Reduced or spatially patterned inhibition, tonic GABAergic conductances^68^, and neuromodulatory states could all modify the local dendritic voltage preceding a CS and thereby alter subsequent Ca^2+^ recruitment^69^. Simultaneous dendritic voltage imaging with glutamate^70^ or GABA^71^ sensors would provide a direct way to identify the synaptic signals that establish these branch-specific pre-spike depolarizations.

The CS Ca^2+^ signal is larger than the SS Ca^2+^ signal throughout the dendritic tree (**Fig. 2b**). This widespread Ca^2+^ elevation may evoke a variety of downstream signaling pathways in different compartments (e.g. soma vs. basal dendrites), plausibly mediating nonlocal plasticity effects. At the same time, transient, apical branch-specific depolarization (plausibly from synaptic excitation) is selectively tagged when it contributes to successful bursting output, providing a high-fidelity mechanism for associative, branch-resolved learning. This view is consistent with theories in which dendritic compartments supply local instructive signals that enable credit assignment across synapses under a global somatic output^57,72,73^. Small bursts (≤3 spikes) failed to recruit distal apical Ca^2+^, consistent with a multiplet plasticity rule^74^.

Place-field (PF) firing showed higher CS prevalence and stronger distal apical recruitment compared to non-PF firing. This difference manifested as sharper spatial tuning of distal dendrites compared to somas. Remarkably, these context-dependent shifts in activity did not affect the empirical V-Ca^2+^ relation. Rather, the elevated apical Ca^2+^ signals in the center of place fields arose from conjunction of apical depolarization and somatic firing, under a constant V-Ca^2+^ mapping.

We suggest that these apical Ca^2+^ signals may play a role in stabilizing or sharpening PF structure. Testing this hypothesis will require longitudinal measurements that link dendritic voltage and Ca^2+^ dynamics to changes in synaptic strength or place-field properties.

The question remains whether our key finding, a compact and deterministic V-Ca^2+^ coupling function, generalizes to other pyramidal neurons such as those in CA1 or the neocortex, and to other brain states or behaviors. CA2 is notably distinct: it is resistant to many forms of long-term potentiation (LTP)^75–77^ and plays a specialized role in social memory^78^, which requires high stability. It is striking that the conjunctive Ca^2+^ mechanism we identified is localized to the apical dendrites, which is the region of CA2 with the strongest evidence for activity-dependent plasticity^34^. It is possible that the rarity of isolated dendritic Ca^2+^ events is a specific evolutionary adaptation of CA2 to preserve stable representations of social identity. In contrast, CA1 and CA3 neurons, which are more plastic, may operate in a high-plasticity regime where isolated Ca^2+^ spikes are more frequent, as has been reported recently^28,30,79^.

The optical and molecular tools introduced here are well suited to address these questions, by probing the structures of dendritic excitations across brain regions, cell types, and behavioral settings. The PISCO microscopy system could also be applied to study other fluorescence signals in three-dimensional samples where optical sectioning and high data-rate are required, such as glutamate^70^, GABA^71^, neuromodulators^80^, or other intracellular signaling molecules^81^. Such measurements could further illuminate the rich interplay of electrical and chemical signaling in neurons.

## Acknowledgements

We thank Andrew Preecha and Camila Bodden for technical assistance. We thank Liam Paninski, Benjamin Antin, Utku Ferah, Chase King, Amol Pasarkar, Ahmed Abdelfattah, and Eric Salter for helpful discussions. XW acknowledges support from an American Heart Association (AHA) postdoctoral fellowship. BHL acknowledges support from National Research Foundation of Korea (NRF) grant RS-2023-00248959. LDL was supported by the Howard Hughes Medical Institute. This work was supported by a Vannevar Bush Faculty Fellowship N00014-18-1-2859, the Harvard Brain Initiative, NIH grants R01-NS126043, R01-NS133755, and R01-MH117042, and Chan Zuckerberg Initiative Dynamic Imaging Grant 2023-321177.

## Author contributions

XW designed the project, developed the optical system, performed animal surgeries and imaging experiments, analyzed data, and wrote the manuscript. BHL developed the genetic construct, optimized the protocol for virus injection and prism implantation surgeries, built the VR system, performed imaging experiments, and analyzed data. PP developed the genetic construct and helped with the experimental design. JDW-C helped with optical system design and data analysis. JX performed histology experiments. SEP and LDL synthesized and provided JF608-HTL. AEC supervised the project and wrote the manuscript. All authors commented on the manuscript.

## Data availability

Data are available on the DANDI archive [link to be added].

## Supplementary Video captions

**Supplementary Video 1. Ca^2+^ and voltage ΔF movies and ΔF/F traces.** Top left: Normalized Ca^2+^ ΔF, linearly interpolated to 1 kHz. Bottom left: Normalized voltage ΔF. Top right: Real-time somatic Ca^2+^ ΔF/F waveform. Bottom right: Real-time somatic voltage ΔF/F waveform.

**Supplementary Video 2. Spatial profile of all Ca^2+^ and voltage events in a representative cell.** Top: Ca^2+^; bottom: voltage. Inset at the bottom left of the voltage panel shows the somatic voltage waveform of the corresponding event. Events were sorted from lowest to highest somatic Ca^2+^ ΔF/F.

**Supplementary Video 3. Normalized spatial profile of all Ca^2+^ and voltage events in a representative cell.** Same data and arrangement as Supplementary Video 2. Each event was normalized by the value in the soma, to highlight the changes in spatial profile.

## Methods Microscope system

The structured illumination spinning disk confocal microscope (**Fig. 1a**, **Fig. S1**) used a 488-nm laser (Coherent, Genesis, MX488, 3W) and a 607-nm laser (Opto Engine LLC, MLL-FN-607, 600 mW) for excitation. A half-wave plate on a rotation mount and a polarizing beam splitter were used to adjust the laser power. The two lasers were combined using a dichroic mirror (IDEX, FF506-Di03-25x36) and coupled into an acousto-optic tunable filter (AOTF, Gooch & Housego, TF525-250-6-3-GH18A) for dynamic modulation of the laser intensity at each wavelength. The two lasers were then separated using another dichroic mirror (IDEX, FF506-Di03-25×36). The 488-nm and 607-nm laser beams were separately expanded by 4× and 8× telescope systems, respectively. The two lasers were then sent to two digital micromirror devices (DMDs, Vialux, V- 7000) for patterned illumination. The DMDs were mounted at a 45° angle relative to the ground, so that the diffracted light stayed in the same horizontal plane as the incident light. The patterned light from each DMD was collected by a separate 4f system, each consisting of two 400-mm convex lenses. Only the central diffraction order was collected. The two lasers were then merged using another dichroic mirror (IDEX, FF506-Di03-25×36).

In commercial spinning disk microscopes, illumination enters through a single-mode optical fiber and is internally homogenized and conveyed onto the spinning disk microlens array. To enable projection of patterned illumination onto the microlens array, we replaced a turning mirror in the side of the spinning disk unit (Yokogawa, CSU-X1) with an optical window in a custom mount (Thorlabs, BCP43R; **Fig. S1b**). The patterned illumination entered through the side window and was re-imaged onto the spinning disk microlens array. The microlens array focused the excitation onto the pinhole array.

An infinity-corrected homebuilt microscope comprised a 100-mm tube lens (TL, Thorlabs, TTL100-A) and a 10× objective (Olympus, XLPLN10XSVMP) to achieve an effective magnification of 5.6×. The microscope re-imaged the spinning disk pinhole array onto the sample. The fluorescence emission from the sample was collected by the same objective-TL pair and re- imaged onto the pinhole array, to achieve optical sectioning. The fluorescence was then reflected by a custom dichroic mirror (Chroma, zt488/607tpc, 13×15×0.5 mm) between the pinhole array and the microlens array. After passing through the spinning disk unit, the two-channel emission light was separated by a dichroic mirror (IDEX, FF580-FDi02-t3-25x36). The green (jGCaMP8m) and red (Voltron2-JF608) emission light was cleaned using separate emission filters (Chroma, ET535/70m for green; IDEX, FF01-709/167-25 for red), and re-imaged onto two scientific CMOS cameras (Hamamatsu, ORCA-Fusion, C11440-22CU) using two separate 100-mm lenses (Thorlabs, AC508-100-A-ML).

The voltage-imaging camera row clock (100 kHz, Hsync) was routed to a data acquisition system (DAQ, National Instruments, PCIe-6323) and used as the master clock for imaging experiments. The Luminos software^38^ was used to coordinate the timing and data acquisition during the experiment and to determine the precise spatial mapping of pixels on each DMD to pixels on the respective camera. To synchronize the voltage-imaging camera exposures with the spinning disk rotation, we used the camera frame clock (1 kHz, Vsync) to trigger a high-quality pulse train from a function generator (Tektronix, AFG3102). The output from the function generator was then connected to the CSU_SYNC (Pin No.12 of the PCR68 connector) port of the spinning disk control unit (**Fig. S2a**). It was necessary to use the function generator, since the Vsync signals from the camera had low duty cycle and slow rise time that resulted in unstable rotation speed of the spinning disk. After starting the camera and spinning disk, we waited 2 s before starting data acquisition, to give the spinning disk time to stabilize its rotation speed. The frame clocks from both cameras were sent to the DAQ to precisely align the timing of the two cameras in post- processing.

## Genetic Construct

The reporter construct, pAAV-hSyn-DIO-Voltron2-LR-p2A-jGCaMP8m (Addgene #251709), was constructed using standard cloning techniques and confirmed via sequencing (Plasmidsaurus). This construct was packaged into AAV 2/9 by UNC NeuroTools core. The AAV9.CamKII 0.4.Cre.SV40 was a gift from James M. Wilson (Addgene viral prep #105558-AAV9).

## Vertebrate animal subjects

All experiments were performed in adult C57BL/6J mice (male, 5-7 weeks old, Jackson Laboratory). All procedures involving animals were in accordance with the National Institutes of Health Guide for the Care and Use of Laboratory Animals and were approved by the Harvard University Institutional Animal Care and Use Committee (IACUC). Animals were group-housed on a reverse 12 h light:12 h dark cycle and had access to food and water *ad libitum*, except during training for and execution of VR experiments when the mice were water restricted, as described below.

## Virus injections

The protocol of virus injections was extensively described in Ref. ^47^. In brief, mice were administered dexamethasone (4.8 mg/kg), carprofen (5 mg/kg) and Ethiqa XR (3.25 mg/kg) ∼1h before the surgery via subcutaneous injection. Mice were then anesthetized with isoflurane and positioned in a stereotaxic frame (Stoelting) with a homeothermic heating pad underneath set at 37 °C. Scalp hair was removed using Nair, and the scalp was disinfected with 70% ethanol and iodine. A scalp incision was made to expose the cranial bone. Three ⌀ 0.5-mm burr holes were made at the stereotaxic coordinates of AP: -1.4 mm, -2.1 mm, -2.8 mm; ML: -1.6 mm using a dental drill (Marathon). The glass micropipettes used for injections were prepared using a Sutter P1000 puller. A microinjection pump (World Precision Instruments, Nanoliter 2010) was used for loading and injecting the virus. The virus mixture comprising AAV2/9-hSyn-DIO-Voltron2-LR- p2A-jGCaMP8m (final concentration: 1-2×10^12^ GC/mL) and AAV9.CamKII 0.4.Cre.SV40 (final concentration: 2-4×10^7^ GC/mL) was injected at all three craniotomy sites at four depths per site (DV: -1.3 mm, -1.2 mm, -1.1 mm, and -1 mm) at a speed of 84 nL/min. 40 nL of virus mixture was injected at each depth, resulting in a total volume of 160 nL per site and 480 nL per animal. After the injection, the scalp was sealed back using Vetbond tissue adhesive (3M). The animal was returned to its home cage with a 37 °C heating pad underneath. The animal’s activity was monitored until full recovery from the anesthesia. Carprofen (5 mg/kg) was administrated via subcutaneous injection for 3 days after the surgery.

## Prism implantation

The prism (2.5 mm height with a 1.5×1.5 mm square top surface; 45° surface coated by aluminum; Tower Optical)^43^ was attached to a 5-mm glass coverslip using optical adhesive (Norland, NOA81). The adhesive was cured by 10-min of UV illumination. The protocol of prism implantation was adapted from Ref. ^43^ and was extensively described in Ref. ^47^. Briefly, the prism implantation surgery was performed 1 week after the AAV injections. The initial procedures up to scalp incision were the same as in **Virus Injections** above. After cranial bone exposure, a 0.5×1.5 mm craniotomy was created using a dental drill centered around the 3 virus injection sites. A single- edge diamond knife (Fisher Scientific, 501930985) was slowly inserted (1 mm/min) through the craniotomy to create a vertical incision with a depth of 1.5-2 mm. The diamond knife was extracted at the same speed to minimize bleeding. Afterwards, a ⌀ 3-mm craniotomy surrounding the diamond knife incision was created using a biopsy punch and a dental drill. The dura mater within the craniotomy was carefully removed using a fine forceps and a Bonn Micro Probe (Fine Science Tools), and the exposed brain was covered by sterile saline. The prism was attached to a blunt-end 14-gauge needle with vacuum and aligned with the incision created by the diamond knife. The prism was then slowly lowered (1 mm/min) inside the incision until the top coverslip was in contact with the skull. During the prism insertion, the side interface of the prism was carefully monitored through the top to ensure that the hippocampal structures (including CA1, CA3, and DG) could be clearly seen. The top coverslip was then fixed on the skull using Vetbond tissue adhesive (3M) and dental cement (Parkell, C&B metabond, 242-3200). Last, a stainless steel headplate was attached to the skull using dental cement. The post-operative procedures were the same as the ones described in **Virus Injections** above.

## Virtual reality (VR) system and training

The VR system was adapted from the design by Christopher Harvey lab (https://github.com/HarveyLab/mouseVR) and the VR environment was designed using the ViRMEn software (https://pni.princeton.edu/pni-software-tools/virmen). Mice were put under water restriction for 3 days (1 mL of water per day) and were trained to run on the VR system for water reward. Animals running more than 50 m in 30 min were considered ready for imaging, typically after 10-14 days of training. The details of the VR system hardware and training procedures were extensively described in Ref. ^47^.

## Image acquisition

Mice were allowed to recover for at least 8 weeks from the prism implantation surgery before imaging. The day before the imaging session, we prepared 100 µL of 0.5 mM JF608-HaloTag solution by dissolving 50 nmol of JF608 powder in 10 µL of DMSO, 10 µL of Pluronic F-127 (20% w/v in DMSO), and 80 µL of 1× PBS. The JF608-HaloTag solution was then delivered via retro- orbital injection at least 12 h before imaging. During the imaging session, the animal was head- fixed and positioned on the treadmill of the VR system. After a target neuron was identified, the mouse and VR system were translated to position the neuron at the center of the field of view. A reference image was acquired and used to manually define DMD patterns for targeted illumination. 60-90 mW/mm^2^ 607-nm light and 5-8 mW/mm^2^ 488-nm light were delivered for illumination. The voltage recording was acquired at 1 kHz and the Ca^2+^ recording was acquired at 41.5 Hz. The VR environment during imaging experiments was different from the training environment and was novel to the animals.

After the functional recording, a 25× objective (Olympus, XLPLN25XSVMP2) was put onto the spinning disk confocal microscope to acquire a high-resolution z-stack image of the neuron structure (0.5-µm axial spacing). Multiple z-stacks were stitched together using the Pairwise Stitching plug-in in Fiji ImageJ. The dendrite tracing was performed on the stitched z-stack image using the Simple Neurite Tracer (SNT)^82^ plug-in in Fiji ImageJ.

## Image processing

All data analysis was performed in MATLAB. **Fig. S6** provides an overview of the pipeline.

### Spinning disk sector correction

The Nipkow spinning disk pinhole array comprised twelve nominally identical sectors, i.e. one revolution of the disk provided twelve scans of the field of view. The voltage-imaging camera acquired one image per sector (twelve per disk revolution). We found that slight variations between the twelve sectors led to period-twelve noise in the voltage-imaging data (**Fig. S2**). We corrected this noise in post-processing, as follows. The temporal average of all frames was used as a reference image. The raw data were then separated into twelve groups based on the remainder after dividing the frame index by twelve (e.g. Group 1: Frames 1, 13, 25…; Group 2: Frames 2, 14, 26…). All frames within each group were averaged, and this average image was divided by the reference image to calculate a group-specific correction pattern. This correction pattern was then divided from all the frames in the corresponding group, and the images from all groups were re- interleaved to reconstruct the corrected data.

This artifact did not occur in the Ca^2+^ imaging data because the frame-rate of the Ca^2+^ camera (41.5 Hz) averaged over two complete rotations of the disk for each frame.

### Voltage and Ca^2+^ image registration

We first obtained the time-average images from the voltage and Ca^2+^ recordings as reference images. We manually selected >15 pairs of control points in each channel, using the *cpselect* function in MATLAB. We then used the *fitgeotform2d* function in MATLAB to calculate an affine transformation to map the Ca^2+^ reference image to the voltage reference image. Once the mapping was visually confirmed to be accurate, the transformation was applied to all the frames in the Ca^2+^ recording.

### Voltage signal extraction

We corrected the sample motion using NoRMCorre ^83^. To identify pixels contaminated by blood vessels, we calculated the lag-1 autocorrection image as *C*(*x*, *y*) = ⟨Δ*F*(*x*, *y*, *n*) · Δ*F*(*x*, *y*, *n* + 1)⟩*_n_*, where *n* is the frame index and ⟨·⟩*_n_* is the average over all frames. Pixels contaminated by blood vessels were highlighted in this image and were manually masked^42,47^. We then manually selected ROIs (20-30 µm in length) in the soma and along each dendrite.

To correct for photobleaching, we first did a rough spike detection and defined silent periods without spiking events. We then obtained the average intensity of all pixels for the silent periods and fitted a two-component exponential decay as an estimate of the bleaching baseline. This bleaching baseline was then divided from the entire recording. We removed residual motion by regressing out the motion traces (*x*(*t*), *y*(*t*), *x*^2^(*t*), *y*^2^(*t*), ndd *x*(*t*) · *y*(*t*) provided by the NoRMCorre algorithm) from the movie.

For each ROI, we performed principal component analysis (PCA) to obtain the voltage footprint (**Fig. S6c**) ^42,47^. Usually the first principal component (PC) had a spatial footprint that matched the structure of the dendrite, and a corresponding time trace which clearly represented voltage dynamics. Other PCs showed spatial structures that indicated residual noise, including sample motion, blood-flow artifacts, animal breathing, and shot noise. We corrected for residual baseline drift by subtracting a low-pass-filtered version of the trace (5-s moving average). Last, we divided by the median value of the low-pass-filtered trace to determine raw ΔF/F voltage dynamics at each ROI.

### Voltage spike detection and classification

The raw voltage ΔF/F trace was high-pass-filtered by subtracting a 50-ms moving-median trace. We then used the *findpeaks* function in MATLAB with a threshold z-score of 5 to detect voltage spikes in each ROI. Spikes with ≤ 1 ms difference in peak timing in neighboring ROIs were linked to the same event. Successive spikes with ≤ 25 ms inter-spike interval at the soma were considered part of the same bursting event. Dendritic spikes without a somatic spike within 4 ms were defined as local voltage dSpikes. We manually examined the raw ΔF movies for all the identified voltage dSpikes to ensure classification accuracy.

Complex spikes (CS) were defined as bursts comprising 4 or more spikes, accompanied by a sufficiently large subthreshold depolarization. To obtain the subthreshold voltages, three frames before and after each spike were removed, and the missing data were filled using linear interpolation. The resulting trace was low-pass-filtered with a 20-ms moving average. Subthreshold depolarizations with peak amplitude exceeding a z-score of 5 (∼50% of the spike amplitude) were considered above-threshold for a CS. All the identified CS events were manually examined to ensure classification accuracy.

The first and last frames of a CS were defined as the points where the rising and falling edges of the depolarization bump crossed a z-score of 1.5. For all non-CS bursts, the duration was defined as 5 frames before the first spike to 5 frames after the last spike. For local voltage dSpikes, the duration was defined as the interval between the two local minima around the peak when the voltage was below a z-score of 1.5. The voltage area under the curve (AUC) was calculated by integrating all frames within the duration of each event.

### Ca^2+^ signal extraction

The Ca^2+^ camera exposure was 24-fold slower than the voltage camera exposure, so there was no need to apply the disk sector correction. We first applied the NoRMCorre motion correction and photobleaching correction as described above. The Ca^2+^ recording was spatially registered to the voltage recording, as described above. We then extracted the raw Ca^2+^ intensity traces using the same ROIs as for the voltage. We estimated slow baseline fluctuations using the *smooth* function in MATLAB (“rlowess” smoothing method with a 100-frame, 2.4 second, smoothing window). This method assigns lower weight to outliers and thus preserves the Ca^2+^ transient peaks. We also manually examined the raw Ca^2+^ intensity traces and the fitted smooth baseline from all ROIs to ensure the preservation of Ca^2+^ spikes. The smoothed baseline was subtracted from the raw traces, and division by the median value of the smoothed baseline resulted in the raw Ca^2+^ ΔF/F dynamics at each ROI.

### Ca^2+^ spike detection and classification

We used the *findpeaks* function in MATLAB with a threshold z-score of 3 (∼5% raw ΔF/F) to detect Ca^2+^ spikes. Dendritic Ca^2+^ events without somatic Ca^2+^ within 2 frames were classified as local Ca^2+^ dSpikes. We manually examined the raw ΔF movies for all the identified Ca^2+^ dSpikes to ensure classification accuracy. For each Ca^2+^ spike, we identified the corresponding voltage dynamics.

For each ROI, the amplitudes of Ca^2+^ spikes were calculated by taking the average of the two largest frames around the peak frame. Negative event amplitude was replaced with 0. Sometimes a Ca^2+^ spike happened on top of the tail of the preceding Ca^2+^ spike. In this case, we calculated the ratio between the minimum in the decaying tail of the earlier event and the maximum of the later event. When this ratio was ≥ 0.3, the later event was excluded from analysis. Otherwise, an exponential decay curve was fitted to the decaying tail of the previous event and extrapolated to the time of the peak of the later event. This exponential fit was then subtracted from the later event before calculating event amplitude.

### Calculating “Robust ΔF/F”

For both voltage and Ca^2+^, each ROI may have different levels of background, which affect raw ΔF/F. To make the ΔF/F values comparable across cells and ROIs, we performed robust estimation of ΔF/F, as shown in **Fig. S6d** and elaborated in Refs. ^42,54^. In brief, for each cell, we calculated the spike-triggered-average (STA) movie of all CS events. For voltage, all events were aligned to the first spike inside the CS, and we took the temporal average of the STA movie to obtain a voltage CS ΔF image. For Ca^2+^, all events were aligned to the peak frame, and we averaged 3 frames centered around the peak frame to obtain a Ca^2+^ CS ΔF image. We also calculated F_0_ images for voltage and Ca^2+^ by taking the temporal average of all silent periods (i.e. no voltage or Ca^2+^ spiking events). For each ROI, we made a pixel-by-pixel scatter plot of ΔF vs. F_0_. We calculated a linear regression for this scatter plot, and the slope of the fitted line represented the robust ΔF/F of CS STA for that ROI (i.e. ∂ΔF/∂F_0_). Any spatially homogenous background signal within the ROI only resulted in an offset of the F_0_ values and did not affect the slope. We also took the average of the same time window for the raw ΔF/F traces to obtain the raw ΔF/F of CS STA. The ratio between ∂ΔF/∂F_0_ and raw ΔF/F of CS STA provided a scaling factor for each ROI. This scaling factor was then multiplied to the raw ΔF/F traces to obtain the robust ΔF/F traces. This robust ΔF/F scaling was calculated separately for each voltage and Ca^2+^ trace in each ROI.

## Contour distance extraction

The high-resolution structural image was first converted into a binary cell mask based on the dendrite traces generated with SNT. The functional voltage imaging data and the manually defined subcellular ROIs were then registered to the coordinate system of the structural image. The position of each ROI was defined as the point within the binary cell mask closest to the center of mass of the ROI. The contour distance of each ROI was then defined as the length of the shortest path within the binary mask connecting that ROI to the soma ROI. Apical contour distances were assigned positive values and basal contour distances were assigned negative values. For 2 out of 13 cells included in this study, the soma was outside the prism imaging field-of-view (**Fig. S3**), preventing the calculation of contour distance. These two cells were therefore excluded from analysis involving contour distance.

## Fast spike timing analysis

For each fast dendritic spike paired with a corresponding somatic spike, we applied cubic spline interpolation to the dendritic and somatic voltage traces to estimate spike peak timing with sub- frame precision. Spike delay was defined as *t_dendrite_* – *t_soma_*. This analysis was repeated for all dendrite-soma spike pairs from cells with somatic recordings. The red dashed lines in **Fig. 1j** define the earliest dendritic spike timing expected for bAP propagation, allowing a time tolerance of *Δt_max_* = 1 ms (one imaging frame). The boundaries were calculated using propagation velocities of 0.22 m/s for apical dendrites and 0.27 m/s for basal dendrites, as determined in **Fig. S7d**. Dendritic spikes occurring earlier than these boundaries were classified as putative dAPs.

## Biophysical characteristics of SS back-propagation

For each cell, we obtained a SS STA voltage trace for all ROIs. We then did a cubic spline interpolation to find the amplitude and timing of the peak in each ROI. For Ca^2+^, we calculated the average amplitude of all SS-induced Ca^2+^ events for each ROI. Within each cell, the peak amplitudes as a function of contour distance from the soma were fit with a single exponential decay curve. This provided voltage and Ca^2+^ SS decay length constants for each cell. To obtain the voltage back-propagation speed, we plotted the SS voltage delay vs. contour distance for all ROIs. We then fitted a line for ROIs ≤ 250-µm away from soma, and the inverse of the slope was the propagation speed.

## Behavioral state classification

The animal’s treadmill speed was smoothed with a 100-ms moving average window. Running and resting periods were classified with a threshold of 1 cm/s. Each voltage event was assigned to a behavioral state based on the average animal speed within the duration of the event.

## Firing rate map, Ca^2+^ ΔF/F map, and spatial tuning curve

To calculate the firing rate map in **Fig. 6e**, we divided the 2-m VR track into 50 equally spaced bins (4 cm / bin). The average firing rate was then calculated for each spatial bin in each lap. Each spike in a CS is counted separately. The Ca^2+^ ΔF/F map in **Fig. 6f** was calculated in a similar way, except that the average Ca^2+^ ΔF/F was calculated for each spatial bin in each lap. The spatial tuning curves in **Fig. 6e, f** were then calculated by averaging the corresponding firing rate maps or Ca^2+^ ΔF/F maps across laps.

## Spatial information

The somatic spatial information based on either CS only or all spikes (**Fig. 6h**) was calculated using the Skaggs formula^84^:

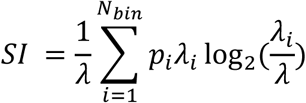

where *λ* is the overall average firing rate, *λ_i_* is the average firing rate at spatial bin *i*, *p_i_* is the probability that the animal is in spatial bin *i*, and *N_bin_* = 50 is the total number of spatial bins.

To compare somatic and distal apical spatial information in the Ca^2+^ channel (**Fig. 6i**), we assigned the amplitude of each Ca^2+^ event to its peak timing and set the values at all other time points to 0, so the analysis was not confounded by the jGCaMP8m decay kinetics. The Ca^2+^ fluorescence- weighted spatial information is calculated as follows^85^:

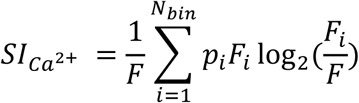

where *F* is the overall average Ca^2+^ ΔF/F, *F_i_* is the average Ca^2+^ ΔF/F at spatial bin *i, p_i_* is the probability that the animal is in spatial bin *i*, and *N_bin_* = 50 is the total number of spatial bins.

## Place cell and place field identification

Place cells and place fields were identified using the criteria described in Ref.^47^: 1) The somatic spatial information was significantly higher than random shuffling; 2) The mean place field firing rate exceeded a defined threshold. The significance of somatic spatial information was evaluated by applying a random circular permutation to the spatial bin index within each lap. A neuron was considered a place cell if its somatic spatial information exceeded the 95^th^ percentile of 1,000 sets of random permutations. For each place cell, the somatic place fields were identified as regions where the average firing rate was greater than 1.5 Hz and the peak firing rate exceeded 5× the average firing rate of the lowest 30% spatial bins. The range of place fields was defined as spatial bins where the average firing rates were larger than 2× the average firing rate of the lowest 30% spatial bins. For place cells, events happening within the range of place fields were classified as PF events, and those outside the range of place fields were classified as non-PF events.

## Place field width

The place field width was quantified using the corresponding spatial tuning curve as follows^26,47^:

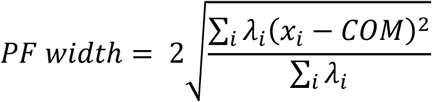

Where *λ_i_* is the average firing rate or Ca^2+^ ΔF/F at spatial bin *i*, *x_i_* is the position of the i^th^ spatial bin in meter, and *COM* is the center of mass of the tuning curve, which is calculated as:

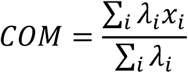

The somatic place field widths based on CS only or all spikes in **Fig.6h** were calculated using the firing rate tuning curve (e.g. **Fig. 6e**), while the somatic vs. dendritic spatial tuning widths in **Fig. 6i** were calculated using the Ca^2+^ ΔF/F tuning curve (e.g. **Fig. 6f**)

## Pre-spike voltage AUC for high vs. low Ca^2+^ events in distal dendrite

We selected ROIs that were ≥ 300-µm from the soma. ROIs within the same dendritic branch were combined. For each CS, we calculated the Ca^2+^ event amplitude in each distal dendrite using the method described above. We calculated the average amplitude of the largest 5 events and used 50% of this value as the threshold to separate high-Ca^2+^ events and low-Ca^2+^ events. Within each category, we identified the corresponding voltage events and aligned them in time to the first spike in the soma. CSs whose first somatic spike occurred < 100 ms after the end of the preceding somatic electrical event were excluded. All remaining voltage events within the same category were averaged to obtain a high-Ca^2+^ and a low-Ca^2+^ voltage STA trace for each branch. This procedure was then repeated across all distal branches. The pre-spike voltage AUC was calculated by integrating the -100 to -2 ms time window before the first spike in the soma (**Fig. 5a**).

## Dendrite-soma voltage difference

To estimate direction of apical-soma current flow, we subtracted the somatic voltage ΔF/F signal from the dendritic signal. To enable comparison between subcellular regions we used the “Robust ΔF/F” method described under **Image processing, “Calculating Robust ΔF/F”**. Positive values indicate a larger depolarization in the dendritic compartment relative to the soma and were interpreted as apical-to-soma current drive; negative values indicate the opposite. The dendrite – soma voltage AUC was calculated by integrating the -1 to 150 ms time window after the first somatic spike (**Fig. 5d**).

## Biophysics-inspired model of voltage-Ca^2+^ coupling

We estimated the value of *E*_Ca2+_ (0.45 in the unit of robust ΔF/F) from the Ca^2+^ equilibrium potential (132 mV with a 2×10^4^-fold difference in extracellular vs. intracellular [Ca^2+^]) and the voltage sensitivity calibration curve of Voltron2-JF608 in Ref. ^41^. The value of *q* (60 in the unit of reciprocal robust ΔF/F) was estimated based on the activation curves of voltage-gated Ca^2+^ channels^58,59,86,87^ and the same calibration curve of Voltron2-JF608. The values of *E*_Ca2+_ and *q* were fixed across all cells and all ROIs. For each ROI, all pairs of voltage event traces and Ca^2+^ event amplitude were used to fit the scaling factor *A* and the activation threshold *V*_0_ by using the *lsqcurvefit* function in MATLAB. A lower bound of 0 was set for *V*_0_ fitting. The same fitted parameters were used without refitting when events were divided into running vs. resting and PF vs. non-PF groups (**Fig. 6b, Fig. S16b**).

## Histology

Mice were transcardially perfused with 1× PBS followed by 4% paraformaldehyde (PFA). The brain was dissected from the skull and post-fixed in 4% PFA at 4 °C for 24 h. 40 µm coronal sections were collected using a vibratome (Leica VT1200S). The brain slices were then rinsed at room temperature with 1× PBS three times for 10 min each, and then blocked for 1 h at room temperature using a blocking solution consisting of 5% bovine serum albumin (BSA) and 1% Triton X-100 in 1× PBS. The brain slices were then incubated with a rabbit anti-GFAP primary antibody (1:500 dilution, Abcam, ab7260) with 1% BSA and 1% Triton X-100 in 1× PBS for 24 h at room temperature. After primary antibody incubation, the brain slices were washed three times with 10 min each at room temperature, with 1% Triton X-100 in 1× PBS. The slices were then incubated with a goat anti-rabbit, Alexa Fluor 488 secondary antibody (1:500 dilution, Invitrogen, A-11008) with 1% BSA and 1% Triton X-100 in 1× PBS for 2 h at room temperature. After secondary antibody incubation, the brain slices were washed in 1× PBS with 1% Triton X-100 three times for 10 min each at room temperature. Afterwards, the slices were mounted onto a glass slide with an antifade mounting medium (Vector Laboratories, H-1900-2). The mounted slides were imaged using a Zeiss Axio Scan.Z1 epifluorescence microscope.

