## Supplementary Information for "Dendrite-wide voltage and Ca^2+^ imaging reveals structures of dendritic excitations during behavior"

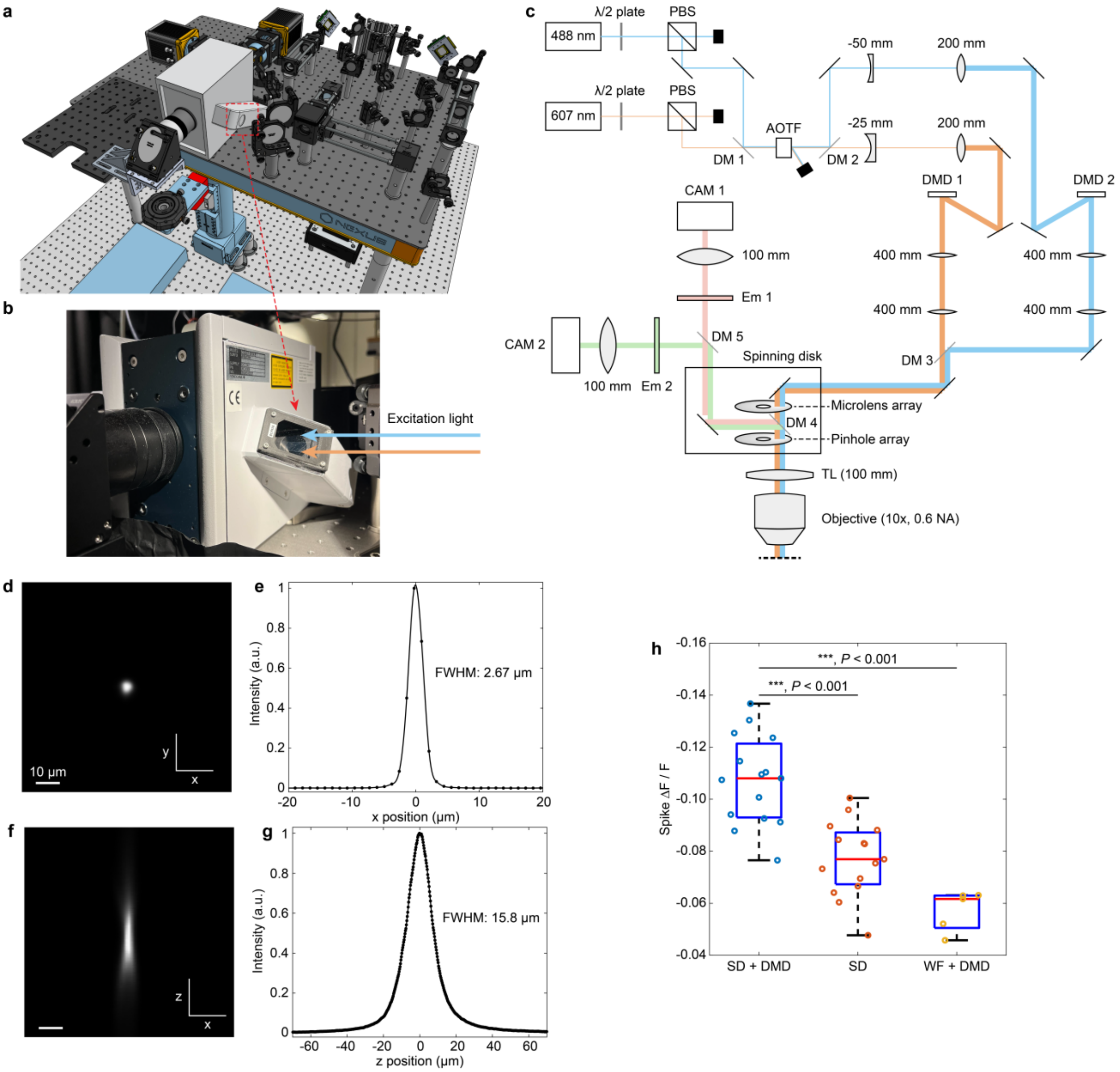


**Fig. S1. Design and characterization of the PISCO system. a,** Overview of the PISCO system. The CAD file is available as **Supplemental File 1**. **b,** A mirror in the side of the Yokogawa CSU-X1 spinning disk system was removed to allow projection of DMD-patterned excitation light directly onto the spinning disk microlens array (**Methods**). **c,** Layout of the optical system. λ/2 plate: half-wave plate; PBS: polarizing beam splitter; DM: dichroic mirror; AOTF: acousto-optic tunable filter; DMD: digital micromirror device; TL: tube lens; NA: numerical aperture; Em: emission filter; CAM: camera. **d-g,** Lateral (**d,e**) and axial (**f,g**) point-spread-function of the optical system. A 2-µm diameter fluorescent bead (ThermoFisher Scientific, F8887) was used as a point source. **h**, Statistics of spike ΔF/F obtained under different optical configurations. 5-15 cells per group. Each point represents the average value from one cell. Most cells were not measurable under wide-field illumination alone. Paired t-test. ***, $P < 0.001$.

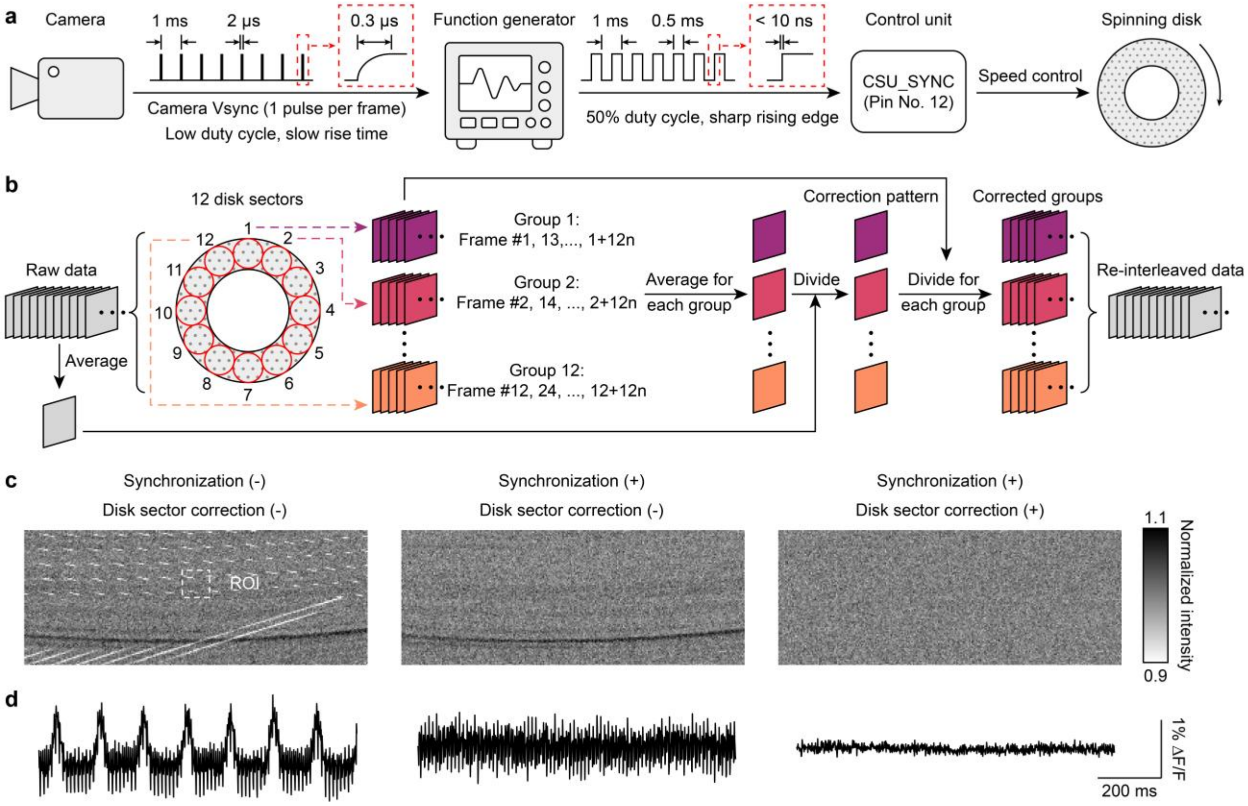


**Fig. S2. Hardware synchronization and computational correction for spinning disk imperfections.** **a**, The camera Vsync frame-clock signal triggered a high-quality pulse train from a function generator. The output of the function generator was connected to CSU_SYNC port of the spinning disk control unit to achieve precise phase-locked synchronization between the camera exposures and the disk rotation. **b**, Algorithm for correcting the slight variation between the 12 spinning disk sectors (**Methods**). **c-d**, Representative images (**c**) and time traces (**d**) acquired without hardware synchronization or disk sector correction (left), with hardware synchronization but no disk sector correction (middle), and with both hardware synchronization and disk sector correction (right).

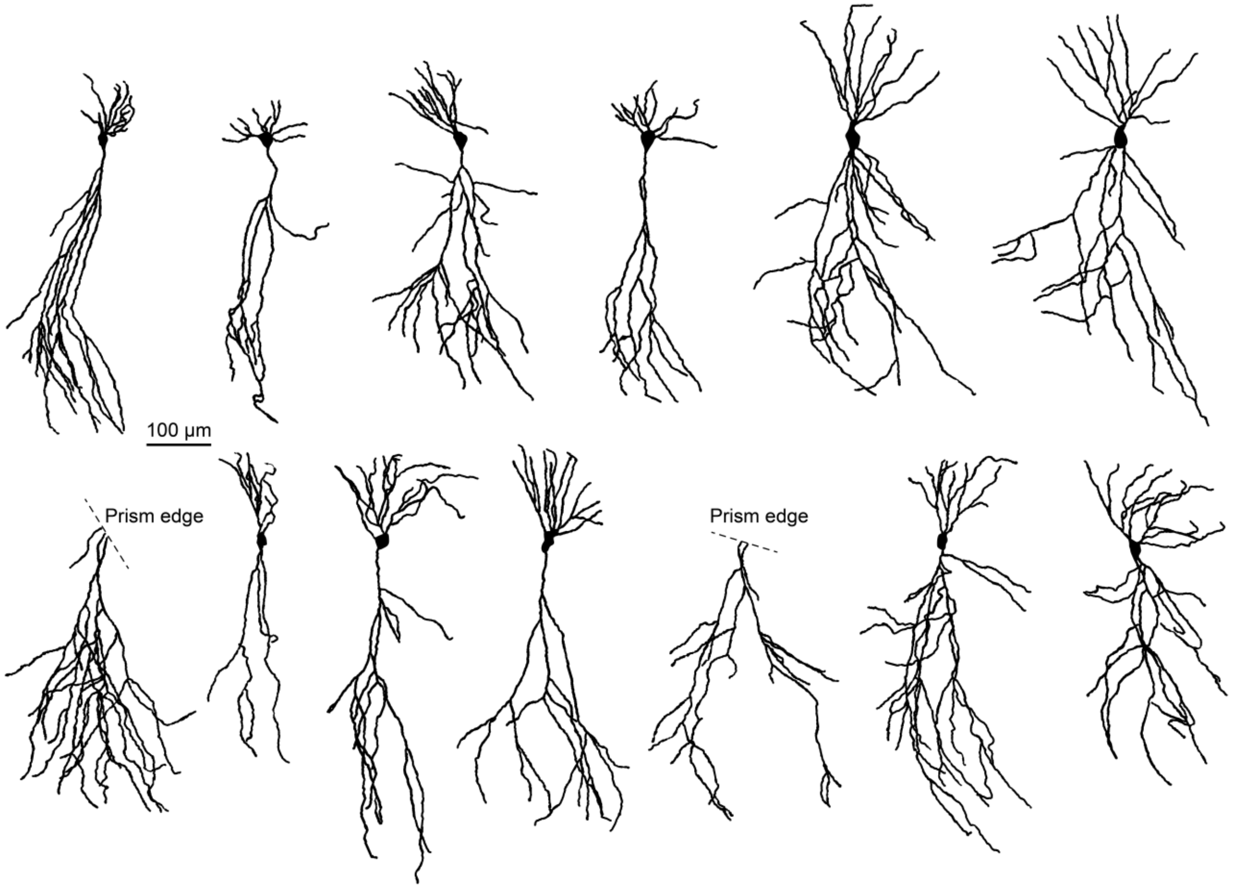


**Fig. S3. Structures of all imaged neurons.** The structures were determined from spinning disk confocal z-stacks, tiled across the entire visible dendritic arbor. The dendrite tracing was performed semiautomatically using the Simple Neurite Tracer (SNT)[1] plug-in in Fiji ImageJ (**Methods**).

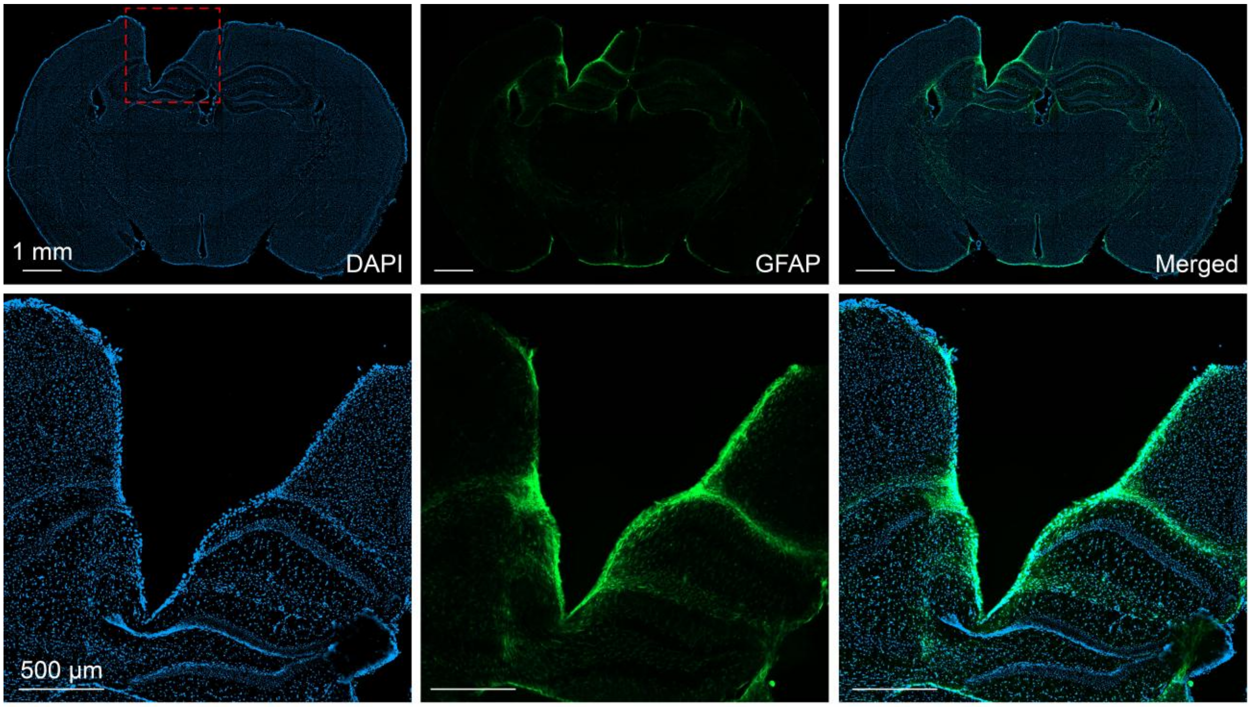


**Fig. S4. Histology at the prism implantation site.** Confocal images showing the distribution of cell nuclei (DAPI, left) and astrocytes (GFAP, middle). The bottom row shows the zoom-in images of the region highlighted by the dashed red square in the top left panel.

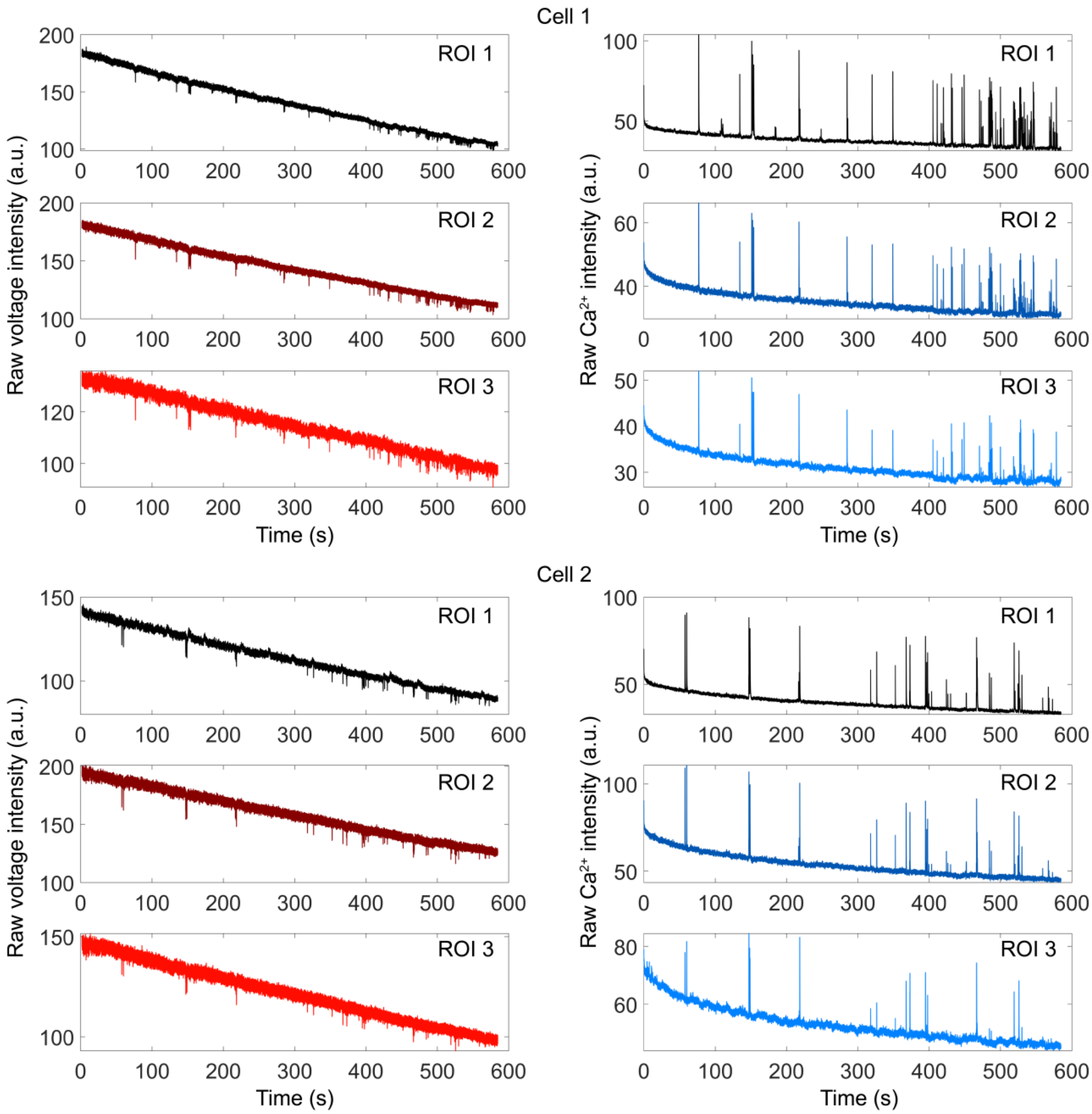


**Fig. S5. Raw data.** Raw 10-min voltage (left) and $Ca^{2+}$ (right) recording traces from two representative cells. ROIs 1-3 for each cell have increasing contour distance from the soma.

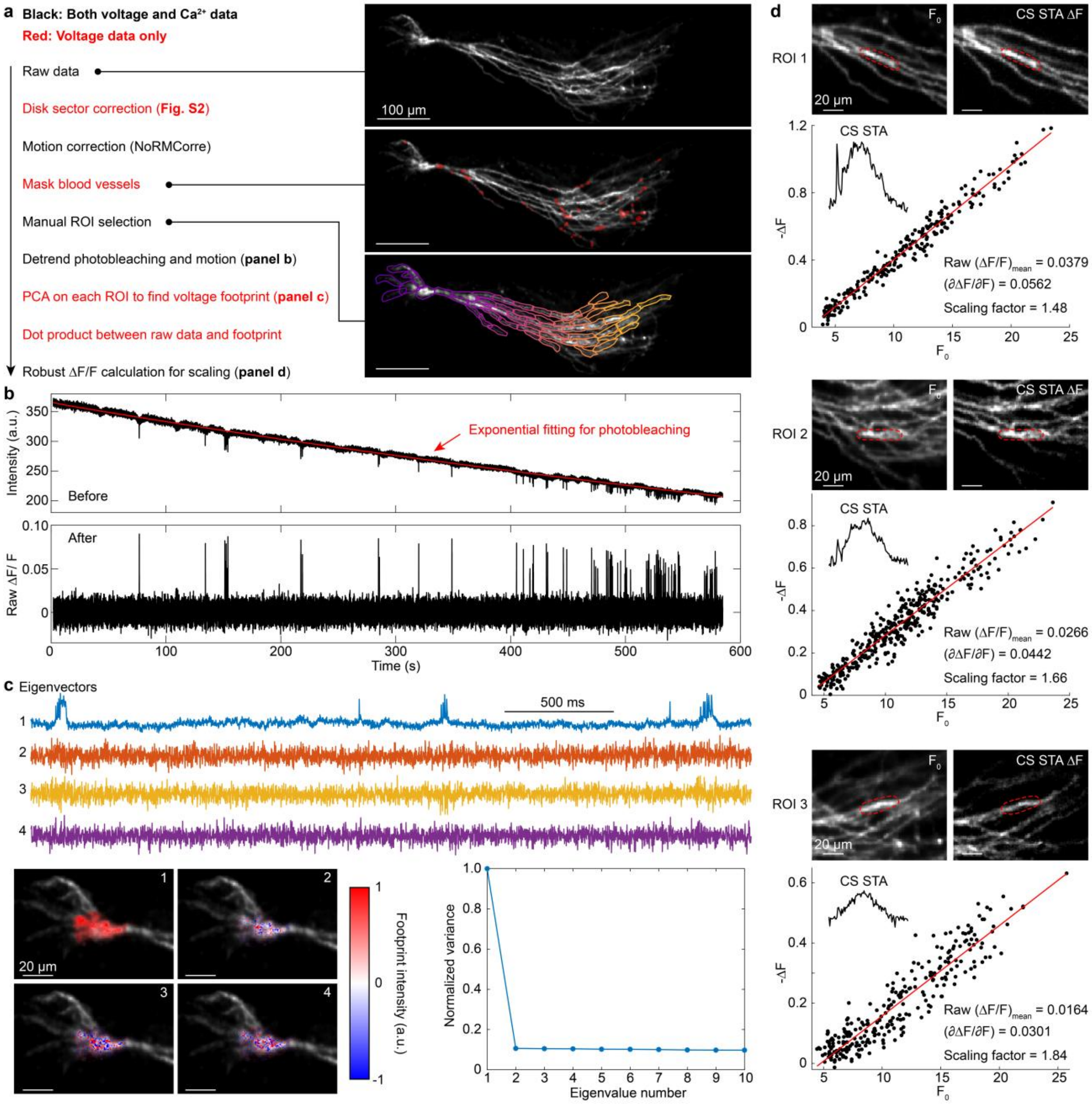


**Fig. S6. Image-processing pipeline. a,** Disk sector correction (as described in **Fig. S2**) was first applied to the raw data, followed by NoRMCorre motion correction[2]. Pixels contaminated by blood vessels were identified and manually masked (**Methods**). Photobleaching was corrected and residual motion was regressed out from the movie. ROIs 20-30 μm in length were then manually selected in the soma and along each dendrite. Principal component analysis (PCA) extracted the spatial footprint and time-trace of the voltage signal in each ROI. Last, to correct for different background values between ROIs, a background-robust ΔF/F analysis[3,4] was applied to each ROI to obtain a scaling factor to convert raw ΔF/F into a signal that was insensitive to background. The three insets on the right show (top) the $F_0$ image of a representative neuron, (middle) pixels contaminated by blood vessels marked in red, and (bottom) manually selected ROIs. **b**, Time traces of the soma ROI before (top) and after (bottom) photobleaching correction. **c**, Top: Time traces of the first four principal components (PCs) from the soma ROI. Bottom left: Footprint of the first four PCs overlaid on the top of a grayscale $F_0$ image. Bottom right: Normalized variance explained by each PC. **d,** $F_0$ images, CS spike-triggered-average (STA) $\Delta F$ images, and pixel-wise scatter plots of $-\Delta F$ vs. $F_0$ in three representative ROIs (outlined by red dashed lines). The insets on the top left of the scatter plots show the waveform of CS STA in each ROI. The slope of the fitted line between $\Delta F$ and $F_0$ is the robust $\Delta F/F$ that is not affected by background signals (i.e. offset in $F_0$)[3,4] (**Methods**).

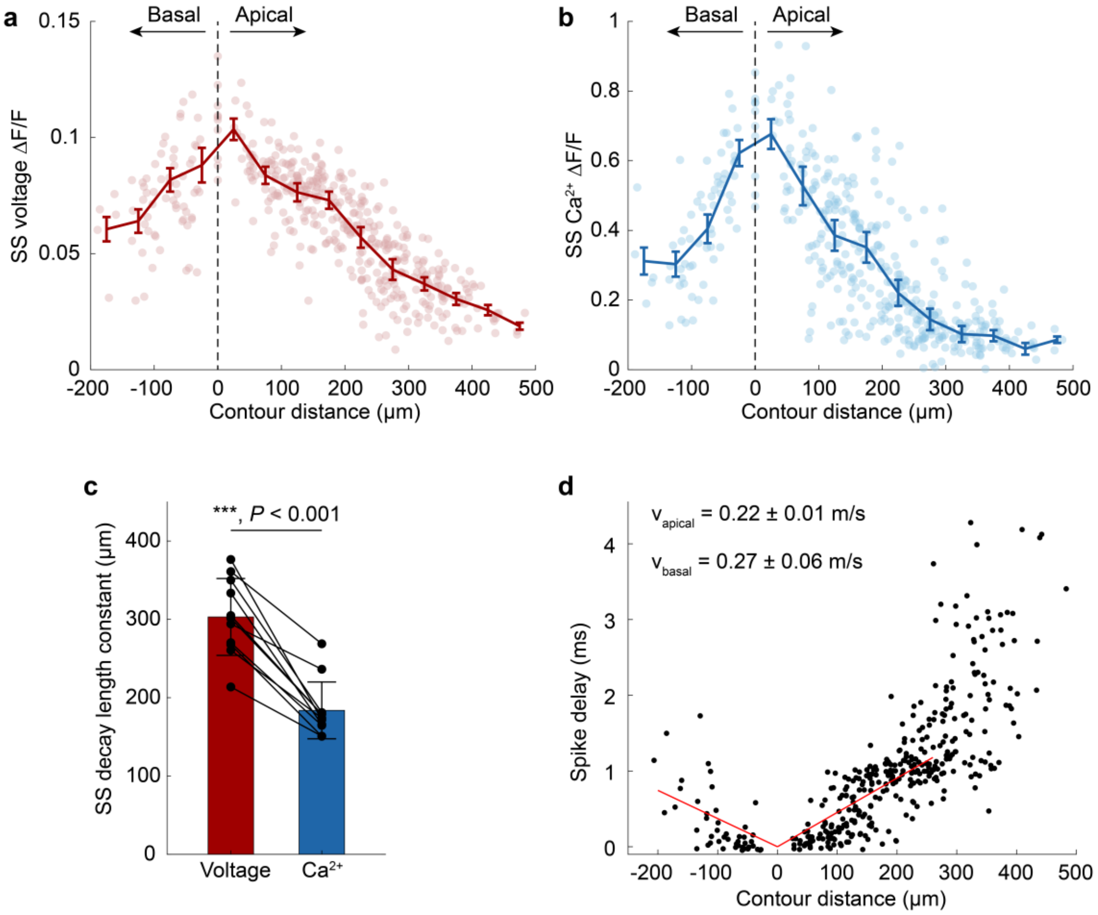


**Fig. S7. Biophysical characteristics of SS back-propagation. a,** Peak voltage and (**b**) $Ca^{2+}$ ΔF/F during SS back-propagation as a function of contour distance from the soma. Data are presented as mean ± s.e.m. Each point represents the average value across all SS events in a subcellular compartment ($n$ = 11 cells, 8 mice). **c,** Exponential decay length constant of voltage and $Ca^{2+}$ during SS back-propagation. Data are presented as mean ± s.d. Each point represents one cell ($n$ = 11 cells, 8 mice). Paired t-test. ***, $P < 0.001$. **d,** Time delay of SS voltage back-propagation as a function of contour distance from the soma. Each point represents the average value across all SS events in a subcellular compartment ($n$ = 11 cells, 8 mice). Solid lines show linear fits for ROIs ≤ 250 μm from the soma. Propagation velocities were calculated as the inverse slopes.

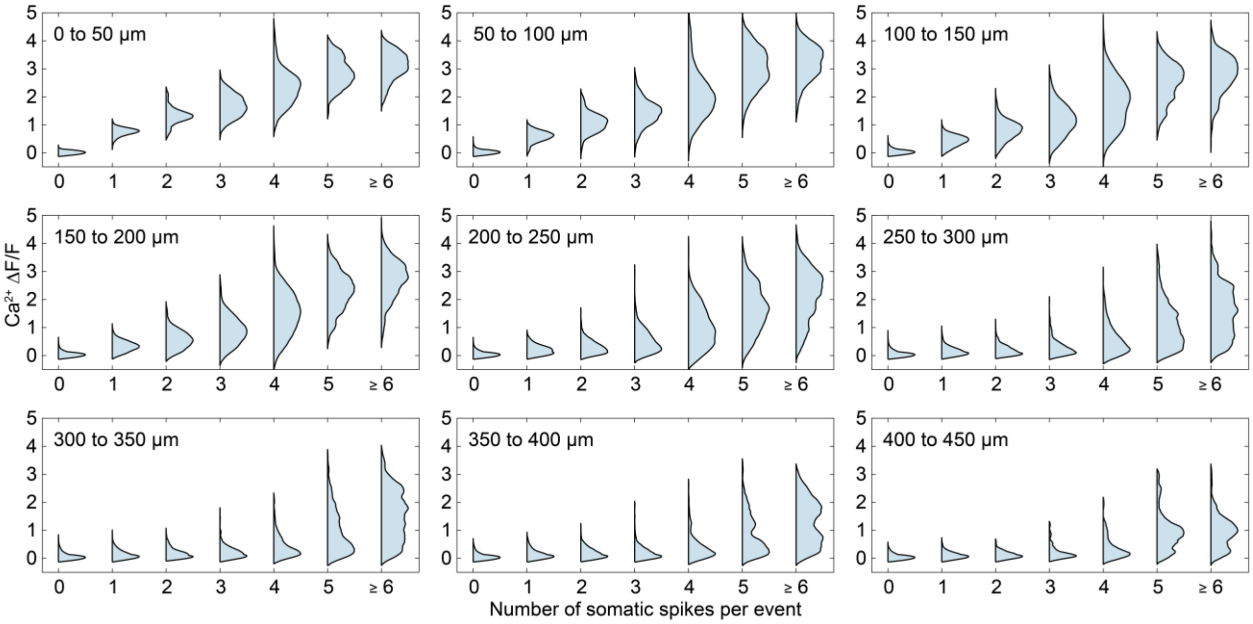


**Fig. S8. Distribution of $Ca^{2+}$ ΔF/F associated with events containing different numbers of somatic voltage spikes.** Data are grouped based on their contour distance from the soma. (*n* = 11 cells, 8 mice, 239 – 1058 events per category). A subset of these plots is also shown in **Fig. 2f**.

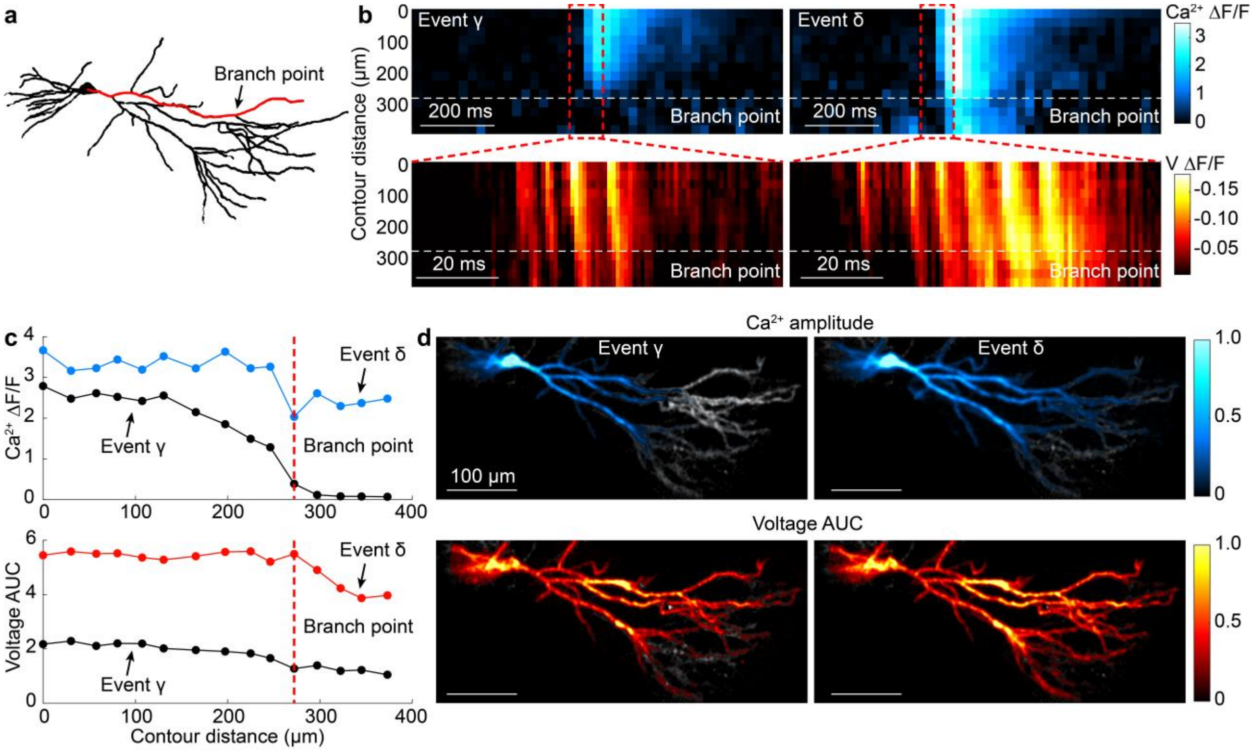


**Fig. S9. Branch point failure. a,** Structure of a neuron with one dendritic path highlighted in red. **b,** Kymographs of $Ca^{2+}$ (top) and voltage (bottom) dynamics for events γ and δ along the path highlighted in (**a**). Note that the $Ca^{2+}$ and voltage kymographs have different timescales. **c,** Line profiles of $Ca^{2+}$ ΔF/F (top) and voltage AUC (bottom) for events γ and δ along the path highlighted in (**a**). The position of the branch point was highlighted with dashed red lines. **d,** Maps of $Ca^{2+}$ ΔF/F (top) and voltage AUC (bottom) for event γ and δ, overlaid on a grayscale structural image of the neuron.

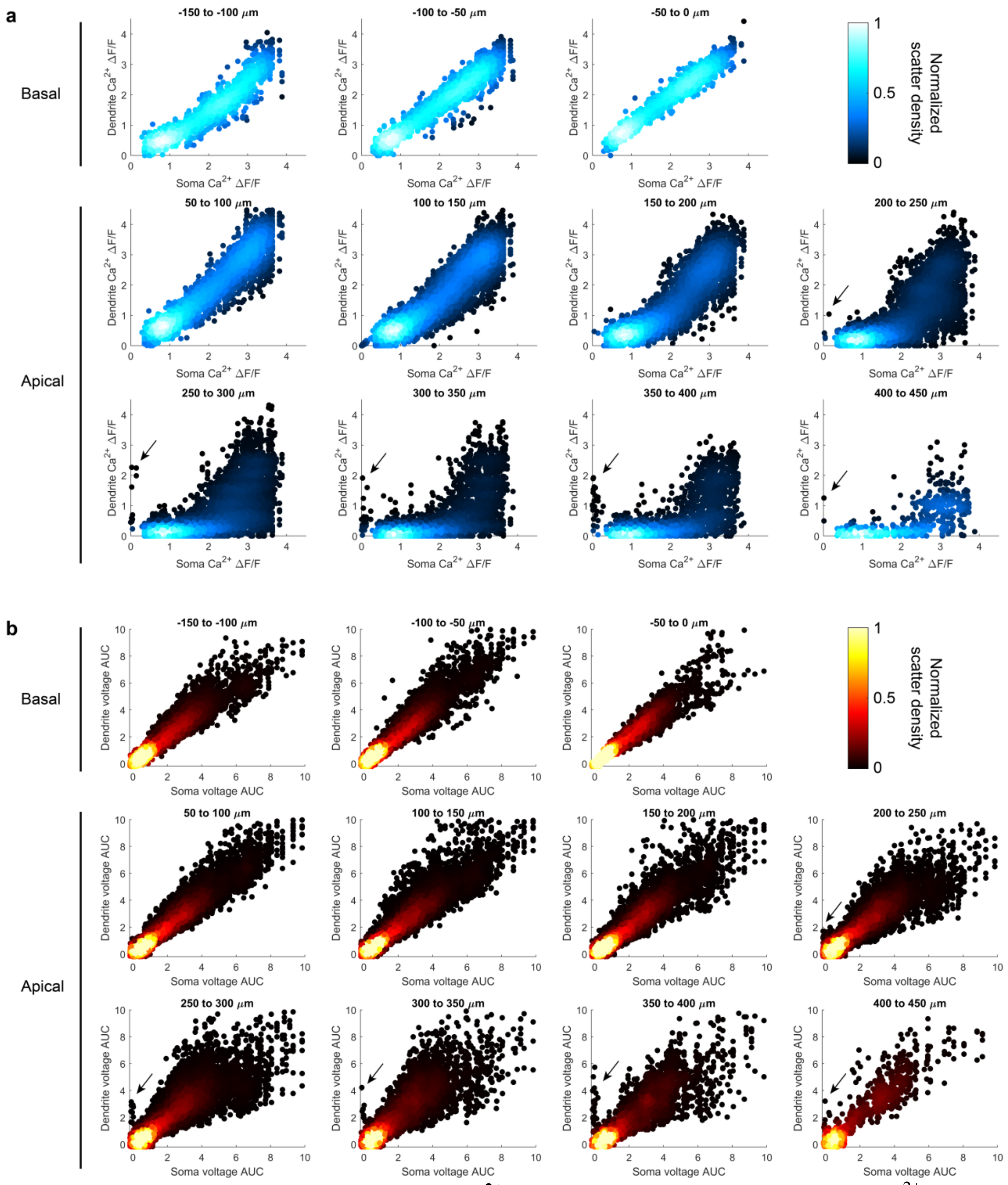


**Fig. S10. Soma-dendrite correlations of $Ca^{2+}$ and voltage signals. a,** Dendritic $Ca^{2+}$ ΔF/F vs. somatic $Ca^{2+}$ ΔF/F at different contour distances from the soma. **b,** Dendritic voltage AUC vs. somatic voltage AUC at different contour distances from the soma. The black arrows indicate local dSpikes. Each point represents one event in a single subcellular compartment (*n* = 11 cells, 8 mice). A subset of these plots is also shown in **Fig. 3h**.

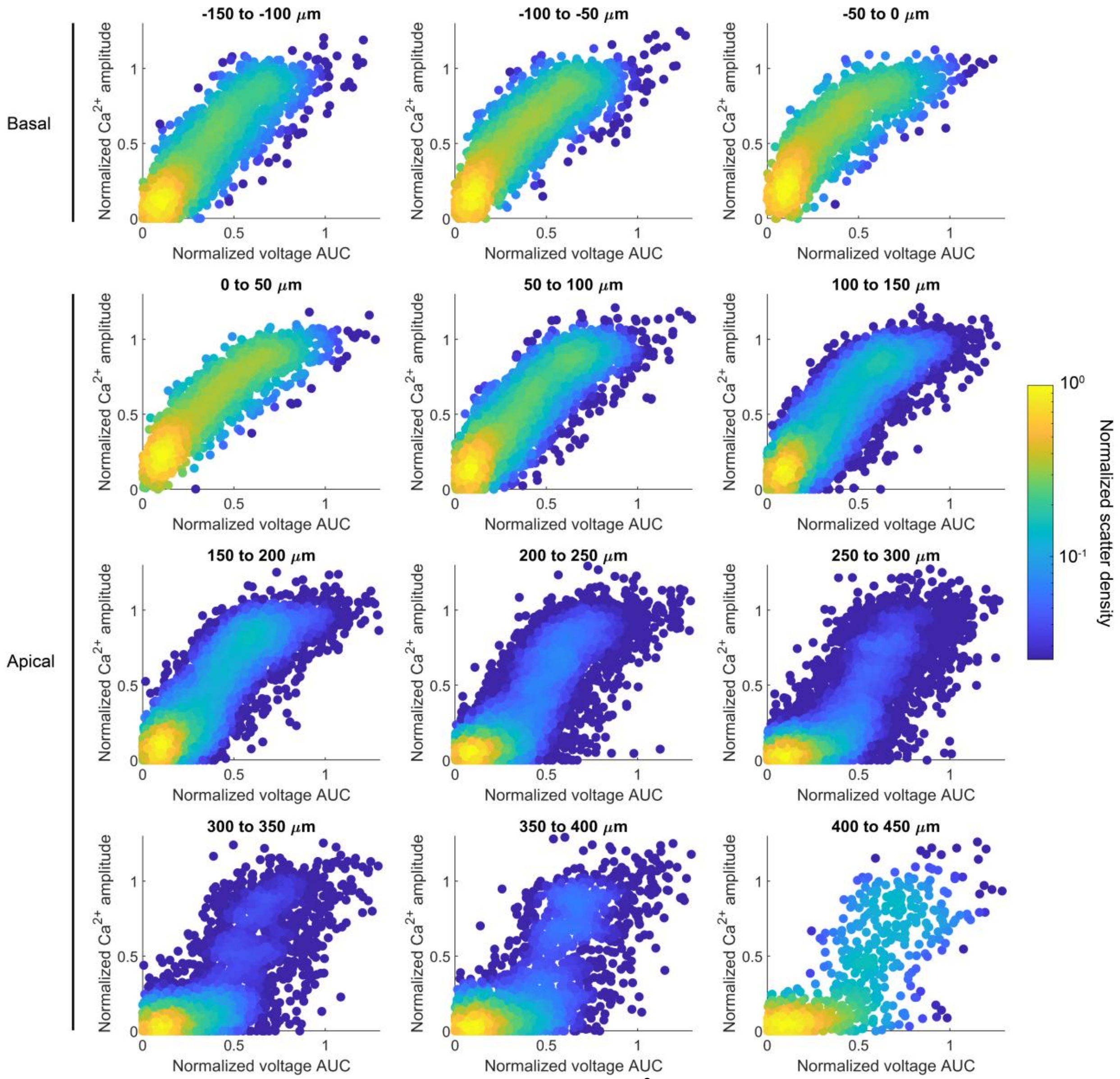


**Fig. S11. Voltage-$Ca^{2+}$ relations.** Normalized local $Ca^{2+}$ amplitude vs. normalized local voltage AUC at different contour distances from the soma. Within each ROI, local voltage AUC and local $Ca^{2+}$ amplitude were independently normalized by the mean of their five largest values. Each point represents one event in a single subcellular compartment (*n* = 11 cells, 8 mice). Five of the plots are also shown in **Fig. 4a**.

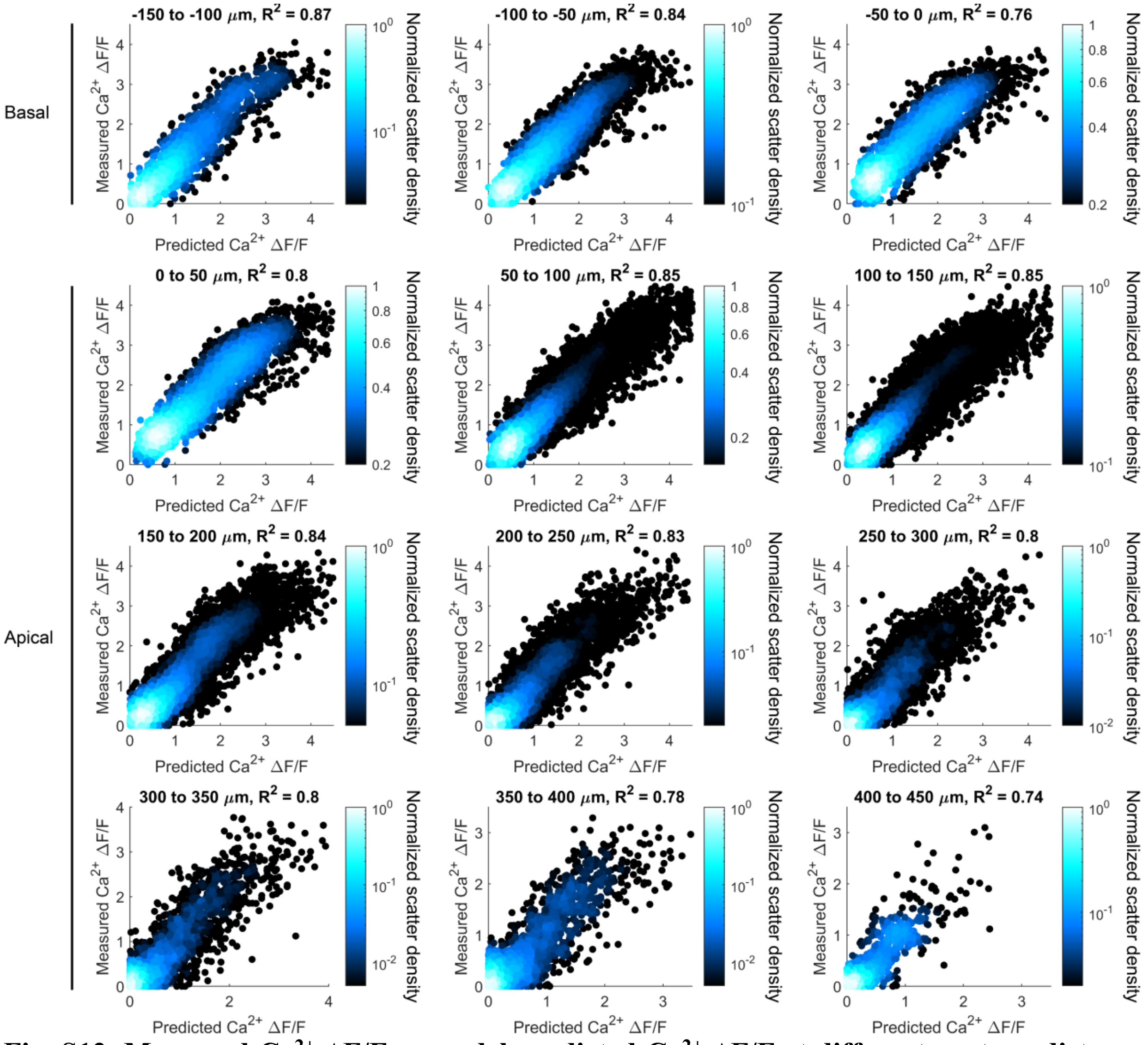


**Fig. S12. Measured $Ca^{2+}$ ΔF/F vs model-predicted $Ca^{2+}$ ΔF/F at different contour distances from the soma.** Each point represents one event in a single subcellular compartment (*n* = 11 cells, 8 mice).

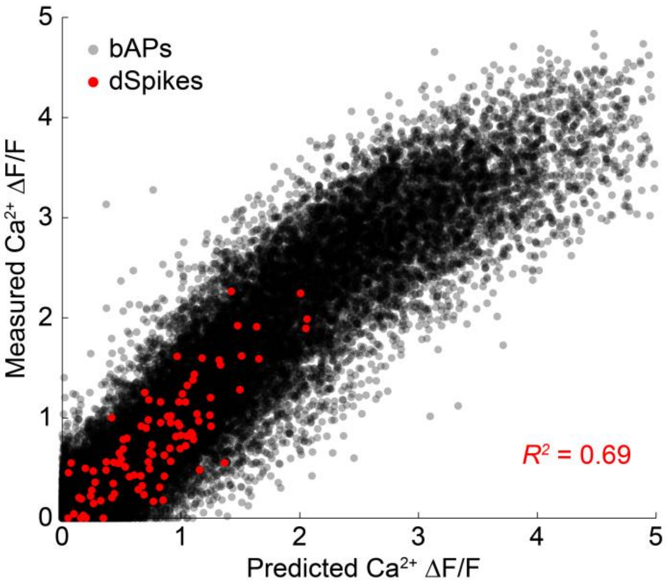


**Fig. S13. Voltage-$Ca^{2+}$ coupling model predicts dSpike $Ca^{2+}$ amplitudes.** Measured $Ca^{2+}$ ΔF/F vs model-predicted $Ca^{2+}$ ΔF/F with the dSpikes highlighted in red. Model parameters were fitted using only bAP-associated events (excluding dSpikes) and were held fixed for the dSpike predictions. Each point represents one event in a single subcellular compartment.

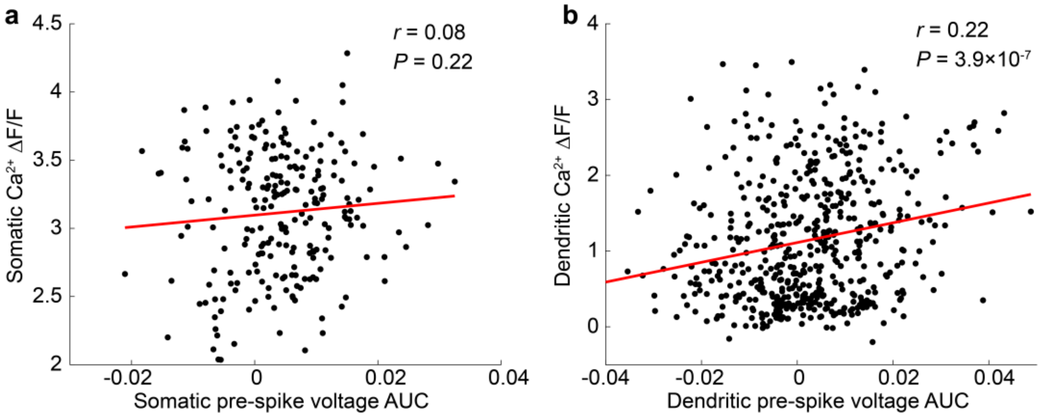


**Fig. S14. Relationship between pre-spike voltage AUC and CS-associated $Ca^{2+}$ amplitude.** $Ca^{2+}$ ΔF/F vs pre-spike voltage AUC (-100 to -2 ms) in the (**a**) soma and (**b**) dendrite. Each point represents one event from a single distal branch (20 distal branches, 8 neurons, 7 animals).

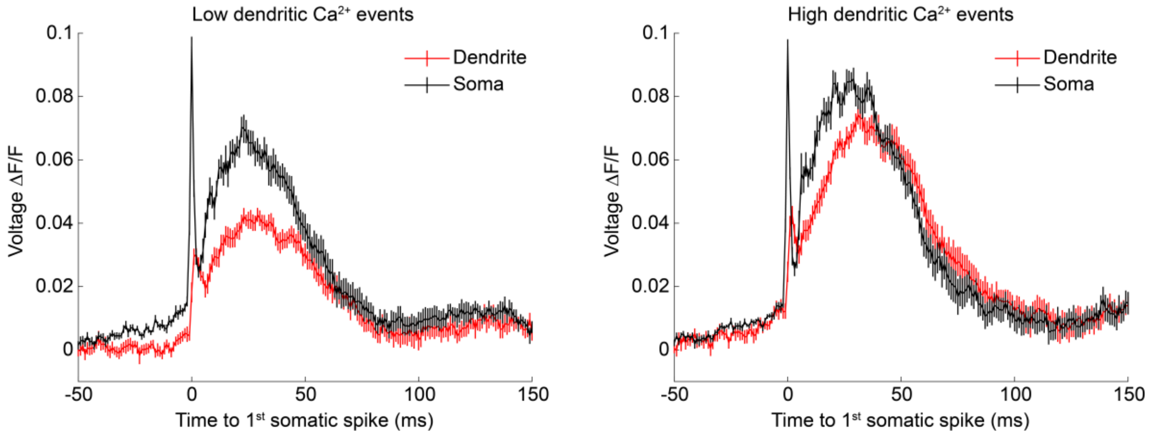


**Fig. S15. Voltage waveforms for high vs. low dendritic $Ca^{2+}$ events.** Average somatic (red) and dendritic (black) voltage waveforms for CS events with low (left) and high (right) dendritic $Ca^{2+}$. Data are presented as mean ± s.e.m. (657 events from 20 distal dendrites in 8 cells). These data were used for calculating the dendrite – soma voltage waveforms in **Fig. 5d** and were also shown in **Fig. 5a** with a different time window.

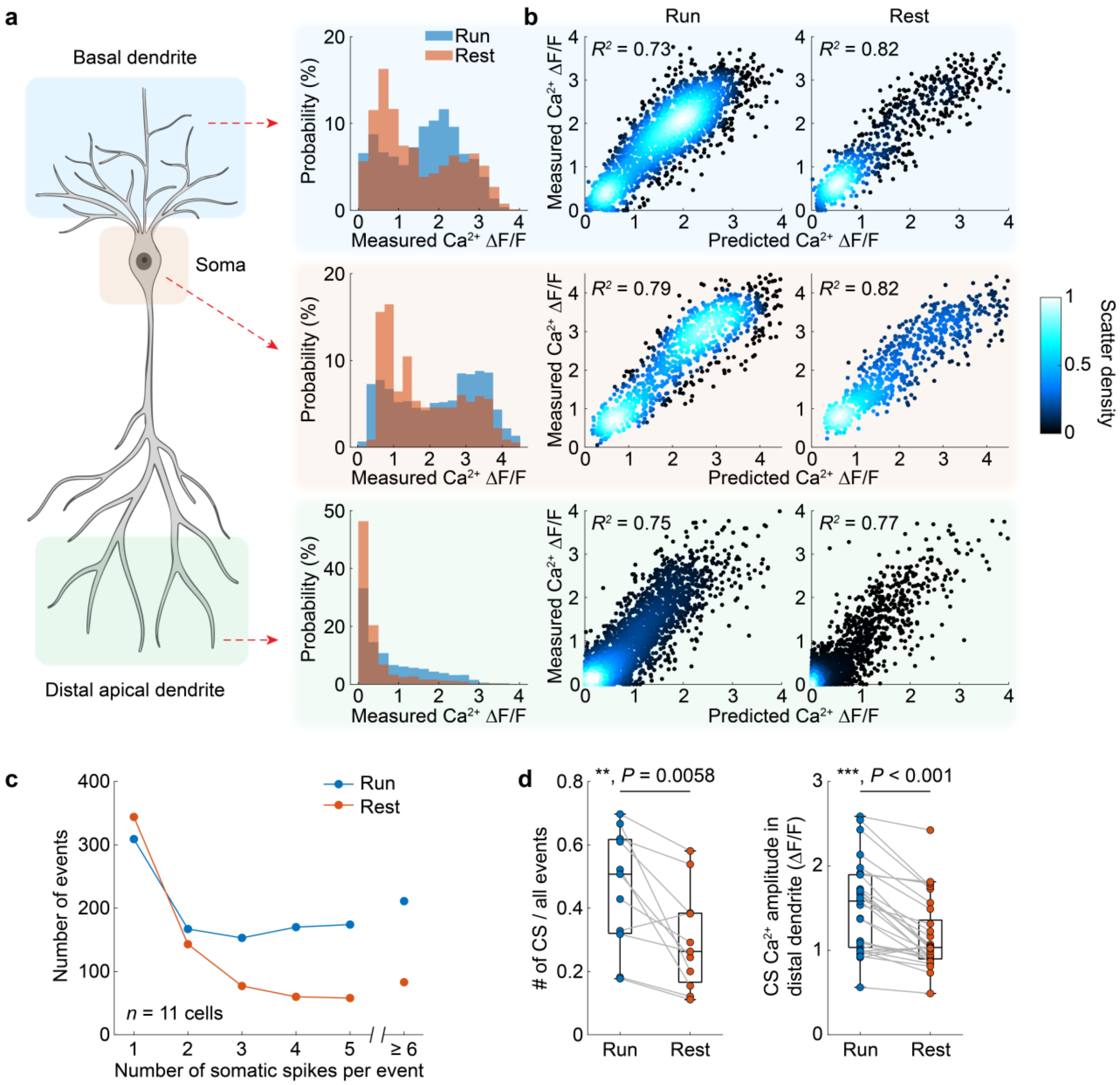


**Fig. S16. Locomotion modulates event prevalence and distal apical $Ca^{2+}$ recruitment. a,** Distribution of measured $Ca^{2+}$ event amplitude during running and resting for basal dendrites (top), soma (middle), and distal apical dendrites (bottom, *n* = 11 cells, 9 mice). **b,** Measured $Ca^{2+}$ ΔF/F vs. model-predicted $Ca^{2+}$ ΔF/F for running and resting events across different subcellular compartments. **c,** Number of events as a function of somatic spike count during running and resting. **d,** Left: Fraction of events classified as CS during running and resting. Each point represents one cell. Right: mean distal apical $Ca^{2+}$ amplitude for CS events. Each point represents one distal branch (*n* = 29 distal branches from 11 cells). Paired t-test.

**Supplementary Table 1. Comparison of high-speed *in vivo* dendritic voltage imaging approaches**

| Ref. | Cell type & behavioral setting | Imaging modality | Voltage imaging rate | Simultaneous functional imaging geometry & dendritic arbor coverage | Simultaneous voltage + $Ca^{2+}$ | Throughput (voxel/sec)† |
|---|---|---|---|---|---|---|
| 5,6 | Cerebellar Purkinje; awake head-fixed | 2P bidirectional line scanning | 2 kHz line rate | Single line scan (~200 µm) of local dendritic segment without soma | Yes | $1\times10^6$ |
| 7 | CA1 pyramidal; spatial navigation | 2P raster scanning | 440 Hz | One local somatic/dendritic ROI at a time (32×32-64 pixels)* | No | $0.45\text{-}0.9 \times10^6$ |
| 8 | Cerebellar Purkinje; awake head-fixed | 2P raster scanning | 440 Hz | 208×24 µm (254×30 pixels) of local dendritic segment without soma | No | $3.4\times10^6$ |
| 9 | CA3 pyramidal; VR spatial navigation | Multi-site 3D acousto-optical 2P line scanning | 1-2 kHz | Sparse multi-site line scans in 3D; 5-14 soma/dendrite ROIs per cell; 1×10-20 µm per ROI | No | $0.15\text{-}0.55 \times10^6$ |
| 10 | CA1 pyramidal; awake behaving | Multi-site 3D acousto-optical 2P line scanning | 1.6 kHz | Sparse multi-site line scans; 1 soma + 3 dendrite ROIs; 2 lines per ROI; 1×6-10 µm per line | Yes | $0.8\text{-}1.3\times10^5$ |
| This study | CA2 pyramidal; VR spatial navigation | PISCO: Patterned illumination spinning disk confocal | 1 kHz | 240×1000 µm contiguous imaging across nearly the entire dendritic arbor (up to 483-µm contour distance) | Yes | $\sim2\times10^8$ |

†Throughput estimated from published scan dimensions and sampling rates.
*Physical size of the imaging ROI not explicitly reported.
